# Associations of early social experience with offspring DNA methylation and later life stress phenotype

**DOI:** 10.1101/2020.08.17.254805

**Authors:** Zachary M. Laubach, Julia R. Greenberg, Julie W. Turner, Tracy Montgomery, Malit O. Pioon, Laura Smale, Raymond Cavalcante, Karthik R. Padmanabhan, Claudia Lalancette, Bridgett vonHoldt, Christopher D. Faulk, Dana C. Dolinoy, Kay E. Holekamp, Wei Perng

## Abstract

In a wild population of spotted hyenas, we tested the hypothesis that maternal care during the first year of life and social connectedness during two periods of early development lead to differences in DNA methylation and fecal glucocorticoid metabolites (fGCMs) later in life. We found that although maternal care and social connectedness during the communal den dependent period were not associated with fGCMs, greater social connectedness after hyenas leave their communal den is associated with lower adult fGCMs. Additionally, more maternal care and social connectedness after leaving the communal den corresponded with higher global (%CCGG) DNA methylation. Finally, we identified multiple DNA methylation biomarkers near genes involved in inflammation that may link maternal care and stress phenotype. Our findings suggest that both maternal care during the first year of life and social connections after leaving the den influence DNA methylation and contribute to a developmentally plastic stress response.

## INTRODUCTION

Early social experiences shape many aspects of an organism’s future phenotype. Starting more than 60 years ago, experiments on laboratory rodents and non-human primates revealed that maternal care and interactions with peers have lasting effects on offspring stress physiology and behavior ^1,2^. In rodents, lower rates of maternal licking and grooming during the offspring’s first ten days caused elevated adult plasma corticosterone in response to external stressors ^3^, and exhibition of fearful behaviors ^4,5^. In addition to maternal care, interactions with group members were protective against an elevated corticosterone response to a standardized stressor in Sprague-Dawley rats ^6^. Similarly, the physiological toll of maternal separation during early life presented as a flat cortisol trajectory in rhesus macaques ^7^. The importance of early social experiences extends to our own species. Studies of children who lived in orphanages found altered Hypothalamic-Pituitary-Adrenal (HPA) activity, including elevated basal cortisol levels that were evident even after adoption ^8^. This body of literature emphasizes the importance of maternal care and early social interactions in the development of a stress phenotype, yet it remains unclear *how* development of the stress phenotype is mediated.

DNA methylation is a mitotically stable epigenetic mark that is responsive to environmental cues and is associated with regulation of gene expression ^9^. It has been championed as a potential biological mechanism linking early social experiences to later stress phenotype. A landmark cross-fostering study in rodents showed that lower rates of maternal licking and grooming corresponded with higher DNA methylation of the promoter region of the hippocampal Glucocorticoid Receptor (GR; *NR3C1*) gene, lower GR RNA expression, and elevated plasma corticosterone among adult offspring ^10^. Subsequent work in rodents demonstrated that variability in maternal care correlated with widespread differences in brain tissue DNA methylation, not only at single promoter regions, but across the entire genome ^11^. In rhesus macaques, maternal vs. peer-led rearing corresponded with genome-wide changes in DNA methylation in both brain tissue and T-cells during adulthood ^12^. Finally, human epidemiological studies show that lower self-reported quality of maternal care corresponds with higher blood leukocyte DNA methylation of brain-derived neurotrophic factor (BDNF) and oxytocin receptor (OXTR) during adulthood ^13^.

Although there is a large comparative literature on this topic, it contains three major lacunae. First, few studies have measured natural variation in the quantity and quality of early social experience. Experimental studies involving maternal separation and peer isolation are routine procedures with captive primates and rodents ^14,15^, but while informative, they do not capture the range of parental care that is likely relevant to offspring development outside of laboratory settings ^3,16,17^. Second, although studies have examined the relationships among social experiences, DNA methylation, and stress phenotype separately, few have measured all three elements in the same population ^10^. Doing so is necessary to explicitly test the hypothesis that DNA methylation represents a mechanistic link between early social experience and future stress phenotype. Third, few studies consider that social experiences vary in form and timing, particularly in wild animal populations where development is subject to natural selection.

Here we examine relationships among early social experience, DNA methylation, and stress phenotype in a long-term field study of wild spotted hyenas (*Crocuta crocuta*). Based on earlier works in primates and rodents, we first hypothesize that unfavorable early social experiences correspond with adverse stress physiology later in life, as indicated by higher glucocorticoid concentrations. Second, we hypothesize that these early social experiences influence patterns of DNA methylation measured in subadult and adult hyenas. Third, we hypothesize that differential DNA methylation is on the causal pathway, and thus, a potential mechanism linking early social experience to future stress physiology.

## METHODS

### Study population

We used behavioral data and biological samples collected between June 1988 and July 2016 by the Mara Hyena Project, an ongoing field study of wild spotted hyenas (*Crocuta Crocuta*) in the Masai Mara National Reserve, Kenya. For each hyena from our study population, we have information on demography, social, and ecological conditions. Using blood samples from immobilized hyenas, we constructed three primary datasets for analyses: global DNA methylation (n = 186), genome-wide DNA methylation (n = 29), and candidate gene DNA methylation (n = 96).

Spotted hyenas are an appropriate species in which to test our hypotheses because they exhibit a wide range of social behaviors, live in large fission-fusion clans that can contain more than 100 individuals, and show a protracted period of maternal care ^18^. Female hyenas reach reproductive maturity at around two years of age and give birth to one or two offspring every 14 to 17 months throughout their lives, starting in their third year of life ^19^. Following a 110-day gestation period, hyena cubs spend the first few weeks of life interacting exclusively with their mothers in a natal den ^20^. During development, hyena offspring rely on their mothers for sustenance and social support until they are at least two years of age ^20^. Young hyenas also socialize with other members of their clan in an earlier phase when hyenas reside in their clan’s communal den – known as the communal den (CD) phase, followed by a later, den-independent (DI) phase when hyenas venture out into their clan’s territory and develop their relationships with other group members ^21,22^.

### Early social experiences

We derived the maternal care variables from focal animal survey (FAS) data collected during observation sessions in which: 1.) mother-offspring pairs were present together for a minimum of five minutes and offspring were less than 13 months old and, 2.) mothers were lactating since our intention was to focus on maternal care received early in life while offspring were dependent on nursing for sustenance. We quantified durations of maternal care behaviors from FAS data ^23^ based on counts of behaviors occurring during each minute of observation in which both the mother and offspring were present together. Behavioral data were collected daily between 0530 – 0900 h and 1700 – 2000 h. We focused on three maternal care behaviors: minutes the mother and cub spent in close proximity (≤1 meter apart), minutes offspring spent latched to their mother’s nipple (nursing), and minutes during which mothers were observed grooming (i.e., licking) their offspring. Information on additional behavioral data processing is in the Supporting Material.

We measured social connectivity by generating association networks among hyenas based on co-occurrences between each hyena and its group members. For each hyena, we constructed association networks during the CD and DI periods. In order to balance our sampling design, we matched the duration of the DI period with that of the CD period. Mean length of both periods was thus was 7.17 ± 0.13 months ^24^.

From each hyena’s association networks during its CD and DI stages, we extracted three metrics that quantify how connected a hyena is with its group members: degree centrality, strength, and betweenness centrality. We focused on these metrics because they measure an individual hyena’s connectedness within its network, which is likely more relevant than overall network structure. A description of social network methods has been published in Turner *et al.* (2018) and is summarized in the Supporting Material.

### DNA methylation: Global (%CCGG), genome-wide, and candidate gene

We quantified global *(%CCGG)* DNA methylation derived from whole blood using the LUminometric Methylation Assay (LUMA) ^25^. Descriptions of the LUMA assay, laboratory procedures, and data cleaning protocol are available in ^26^. The majority of CpG sites assessed via LUMA occur in gene bodies, where they may function in transcription regulation and alternative splicing ^27^, as well as in non-coding regions of the genome, where they may repress repetitive elements ^28^ and enhance chromosome stability ^29^. Therefore, we assumed that lower than average %CCGG methylation is likely disadvantageous to health.

We measured genome-wide DNA methylation at a single nucleotide resolution in whole blood collected from hyenas 11-27 months old. To prepare the multiplexed Enhanced Reduced Representation Bisulfite Sequencing (mERRBS) sample library, we followed the protocol of Garrett-Bakelman et al. ^30^ using 100ng of high quality genomic DNA. Each sample was spiked with 1ng of non-methylated Lambda gDNA to estimate bisulfite conversion efficiency prior to library preparation. Five libraries were pooled per lane of an Illumina HiSeq4000^®^ platform for single-end sequencing with a 50-nucleotide read length. After DNA sequencing, we ran a standard bioinformatic pipeline to clean, align, and call DNA methylation reads. Details on DNA library preparation and the bioinformatics pipeline are in the Supporting Material.

Given the extensive literature on the relevance of the glucocorticoid receptor (GR) gene to both early social experiences and stress phenotypes, we also assessed CpG methylation in the putative GR promoter region of DNA from 96 hyenas. Candidate gene bioinformatics and laboratory methods are in the Supporting Material. We identified CpG sites in the hyena genome that aligned with those in DNA from humans and rats (Figures S2-S6 in the Supporting Material ^31,32^). We calculated CpG site-specific DNA methylation in the putative hyena GR promoter region using pyrosequencing.

### Fecal glucocorticoid metabolites (fGCMs)

Since January 1993, we have opportunistically collected fecal samples any time an individually identified hyena was seen defecating. Fecal samples were mixed and transferred to 2mL cryovials before flash freezing in liquid Nitrogen within 12 h of collection. The samples were then transported from our field site to the U.S. Here, we focus on the stress hormone corticosterone, measured via a validated hormone extraction process and radioimmunoassay developed for our study population ^33,34^.

### Demographic, social experience, and ecological covariates

We considered three categories of potential confounding variables (i.e., those that are associated with the explanatory variables and potential determinants of the dependent variables) and included them in multiple variable models to improve causal inference. Age and sex were demographic confounders; maternal rank, litter size, parity and clan size were social experience confounders; and human disturbance and local prey abundance were ecological confounders; prey abundance was dichotomized by the approximate timing of the annual mass migration of wildebeest and zebra. The Supporting Material provides details on data collection germane to these variables.

### Statistical modeling framework

We analyzed data in four distinct steps, in accordance with our goals. First, we characterized the relationship between early social experience, according to maternal care and social network metrics (explanatory variables) and adult stress phenotype as indicated by adult fGCMs (dependent variable). Second, we examined associations between early social experience and global DNA methylation – a presumed marker of genomic stability and overall health, and between global DNA methylation and fGCMs. Third, given that global DNA methylation may be considered both as a health outcome and as a potential mechanism, we conducted a formal mediation analysis following Baron & Kenny’s method (1986) on a subset of hyenas for which we had data on early social experience, global DNA methylation, and adult corticosterone levels. Fourth, to complement the third objective, we used genome-wide DNA methylation data to identify potential functional biomarkers that might link early life experience with future stress phenotype and that might be assessed as mediators in future analyses.

### Part 1: Associations among early social environment, DNA methylation and adult *fGCMs*

First, we examined each maternal care variable (proportion of time in close proximity, nursing and grooming) and social network metric (degree centrality, strength, and betweenness centrality in the CD and DI stages) as explanatory variables in relation to fGCMs as the dependent variable. In these models, we considered covariates that are associated with the explanatory variable and potential determinants of the dependent variable (confounders), as well as factors that account for variability in either the exposure or the outcome (precision covariates). Offspring sex and level of human disturbance during the year in which offspring were born (low, medium, and high) were included as potential confounders. We also controlled for the following precision covariates: hyena’s age, reproductive state for females (nulliparous, pregnant, lactating, or other), and time of day (A.M. or P.M) when the fecal sample was collected. For these models, we used mixed effects linear regression with a random intercept for hyena ID to account for correlations between repeated measures of fGCMs (**Figure 1**).

**Figure 1.**
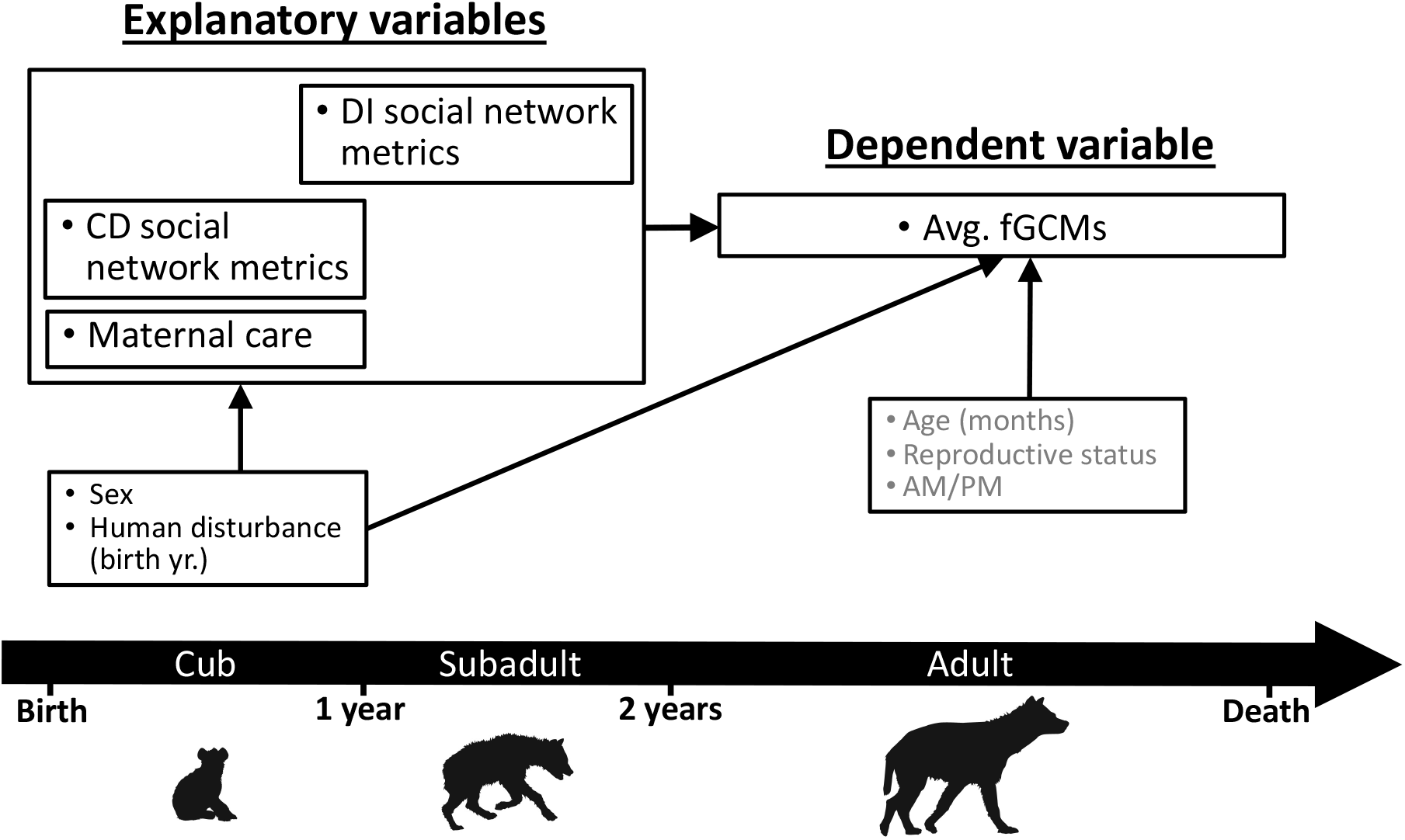
Directed Acyclic Graph (DAG) showing assessment of associations among early life social experience and adult fecal Glucocorticoid Metabolites (fGCMs). Covariates were included based on *a priori* biological knowledge and bivariate analysis (see Supporting Tables S2-S5). Precision covariates (gray text) were assessed when the fecal sample was collected. Variable positions over the timeline roughly corresponded with the timing of assessment.

### Part 2: Associations between early social environment and global DNA methylation, and between global DNA methylation and fGCMs

We modeled the associations between each maternal behavior variable and social network metric as separate explanatory variables with global DNA methylation (%CCGG methylation) as the continuous dependent variable of interest. In all models, we included a random intercept for maternal ID to account for correlations among siblings, and included the covariates depicted in **Figure 2**.

**Figure 2.**
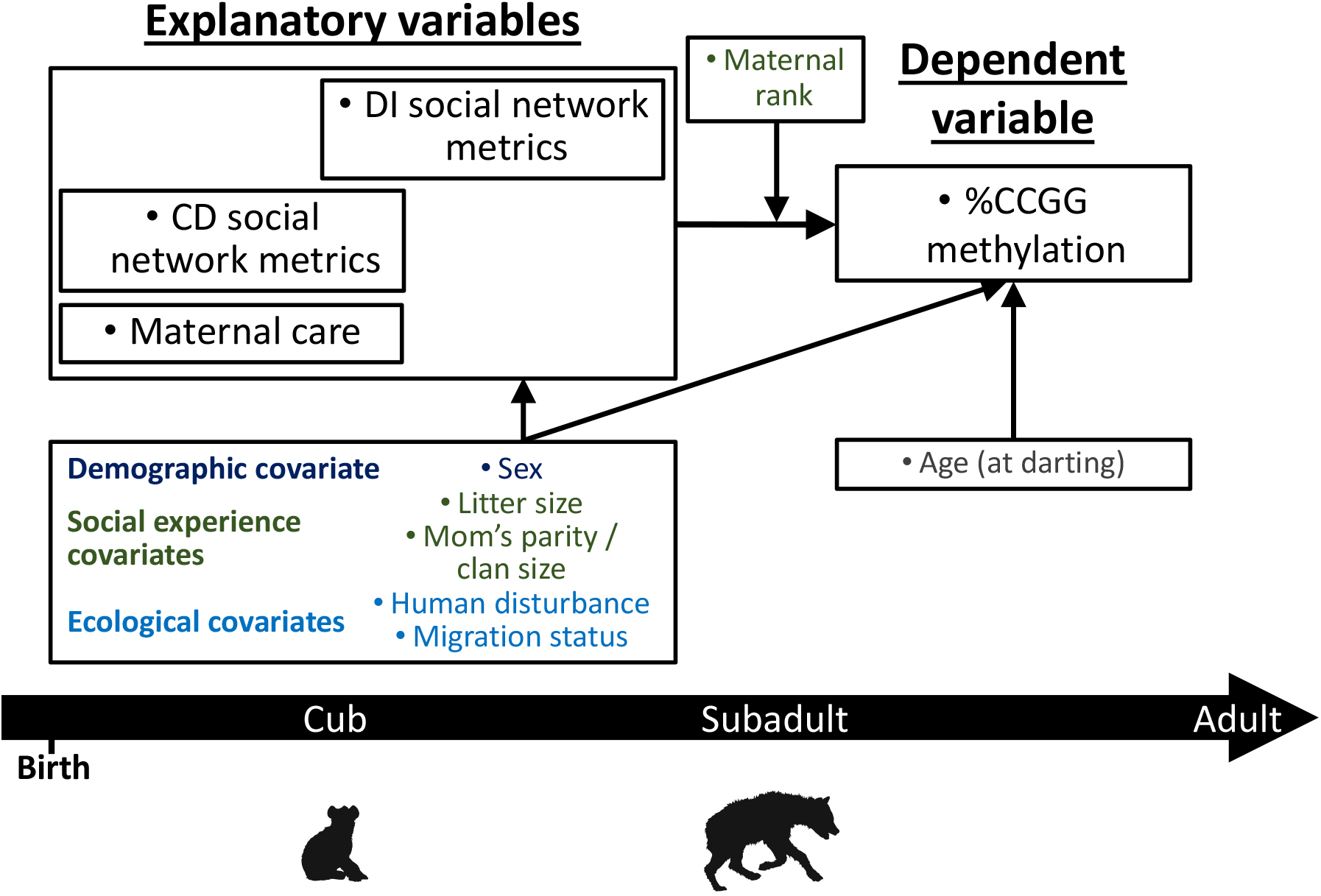
Directed Acyclic Graph (DAG) showing Part 2 of our analysis: assessing associations between early life social experiences and %CCGG methylation. Potential confounding variables were modeled in three different context groups, which included demographic covariates (dark blue), early social experience covariates (dark green), and ecological covariates (light blue). Maternal rank was considered a potential effect modifier of the associations of interest, and age at darting was considered a precision covariate. Variable positions over the timeline roughly corresponded with the timing of assessment.

In multiple variable analysis, we controlled for offspring age in months as a precision covariate because age is a known determinant of DNA methylation in this and other species ^26^. Then, in models where a maternal care behavior was the explanatory variable, we further accounted for three categories of covariates using separate models: demographic covariates (offspring sex); social experience covariates (number of littermates, mother’s parity, and mother’s social rank), ecological covariates (human disturbance and prey abundance on the date offspring were born), and a fully adjusted model that included all covariates. When CD or DI social network metrics were the explanatory variables of interest, we controlled for the same confounders, except that we included clan size rather than the mother’s parity in the social experience covariates model. Given the importance of maternal rank to offspring DNA methylation ^26^, and that maternal rank may mitigate or exacerbate effects of mother-offspring interactions ^36,37^, we tested for statistical interactions with maternal rank in models in which %CCGG methylation was the outcome. We considered α = 0.10 for the interaction term as evidence of effect modification by maternal rank.

We also assessed the relationship between global DNA methylation and adult fGCMs (**Figure 3**). We used a linear mixed model with a random effect for individual ID to account for repeated fGCMs assessments and controlled for sex and level of human disturbance during the year the offspring was born. Precision covariates included the hyena’s age, reproductive state, and time of day when the sample was collected. In all models, we considered α = 0.05 as the threshold for statistical significance, unless otherwise indicated.

**Figure 3.**
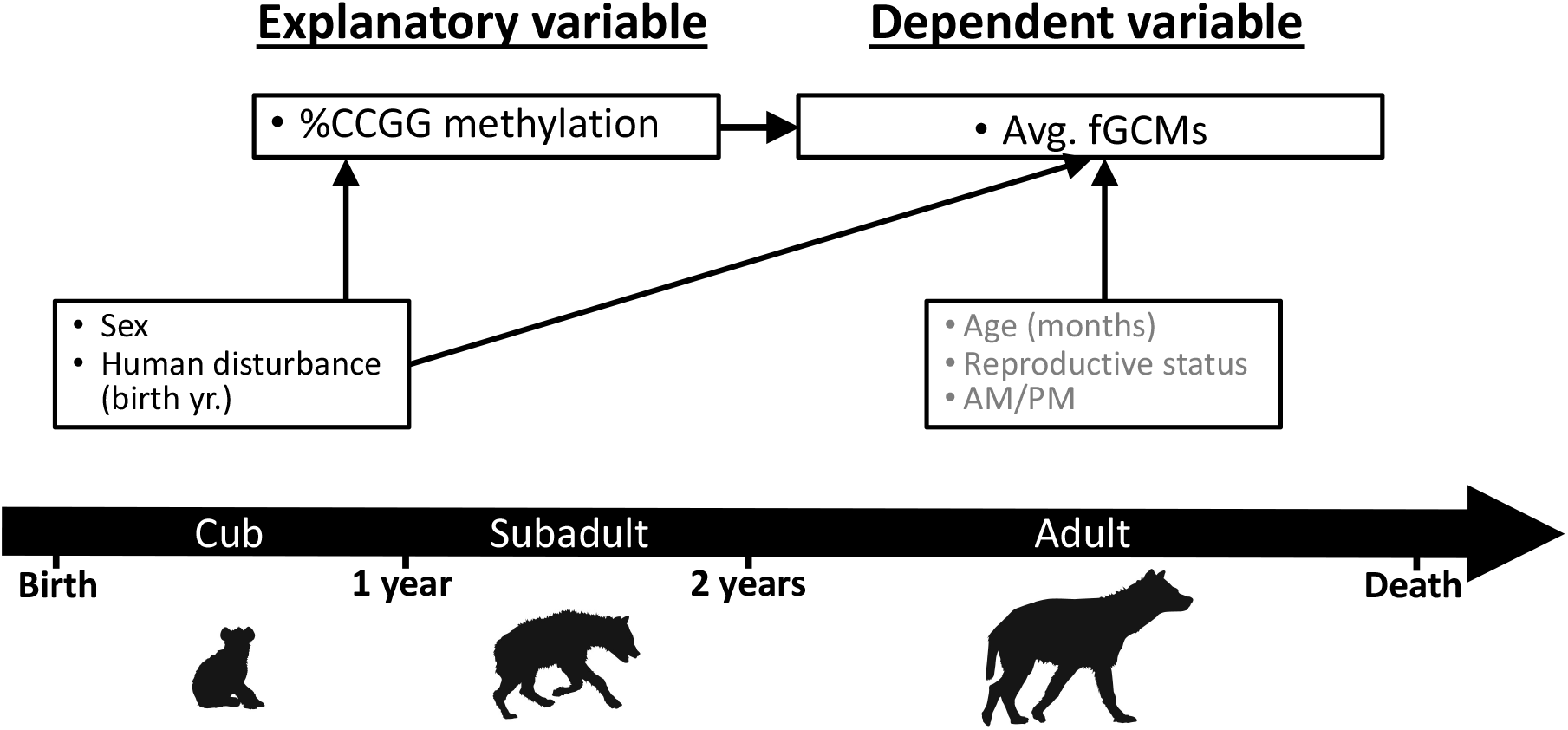
Directed Acyclic Graph (DAG) showing assessment of the associations of %CCGG methylation with adult fecal Glucocorticoid Metabolites (fGCMs). Covariates are included based on *a priori* biological knowledge and bivariate analysis (see Supporting Tables S2-S5.). Precision covariates (gray text) are assessed when the fecal sample was collected. Variable positions over the timeline roughly correspond with the timing of assessment.

One of our aims was to examine associations with DNA methylation of the GR promoter region among 78 hyenas with high quality DNA. However, our assays revealed invariant and near zero percent methylation at six CpG sites, including the putative *NGFI-A* transcription factor binding site, so no additional analyses were performed with respect to this region.

### Part 3: Formal mediation analysis

The third part of our analysis assessed the extent to which maternal care and social network metrics during CD or subadult DI life stages are associated with adult fGCMs before and after adjustment for global DNA methylation (%CCGG) to assess this metric as a mediator in the subsample of hyenas for which we had complete data on early social experience, DNA methylation, and stress phenotype. Following Baron & Kenny’s mediation procedure (1986), we first tested whether maternal care or social network metrics were associated with fGCMs (**Figure 4**). Next, we assessed whether the early social experience variables were associated with %CCGG methylation, and whether %CCGG methylation was associated with fGCMs. Finally, if associations from Steps 1 and 2 were significant, in Step 3 we included %CCGG as an additional covariate in models in which each early social variable was the explanatory variable and fGCMs were the outcome (Step 3; **Figure 4**) and compared estimates of association for each early social variable with vs. without inclusion of the DNA methylation mediator ^35^. We considered evidence for mediation by DNA methylation if inclusion of DNA methylation attenuated the estimate for a given early social metric by >10%.

**Figure 4:**
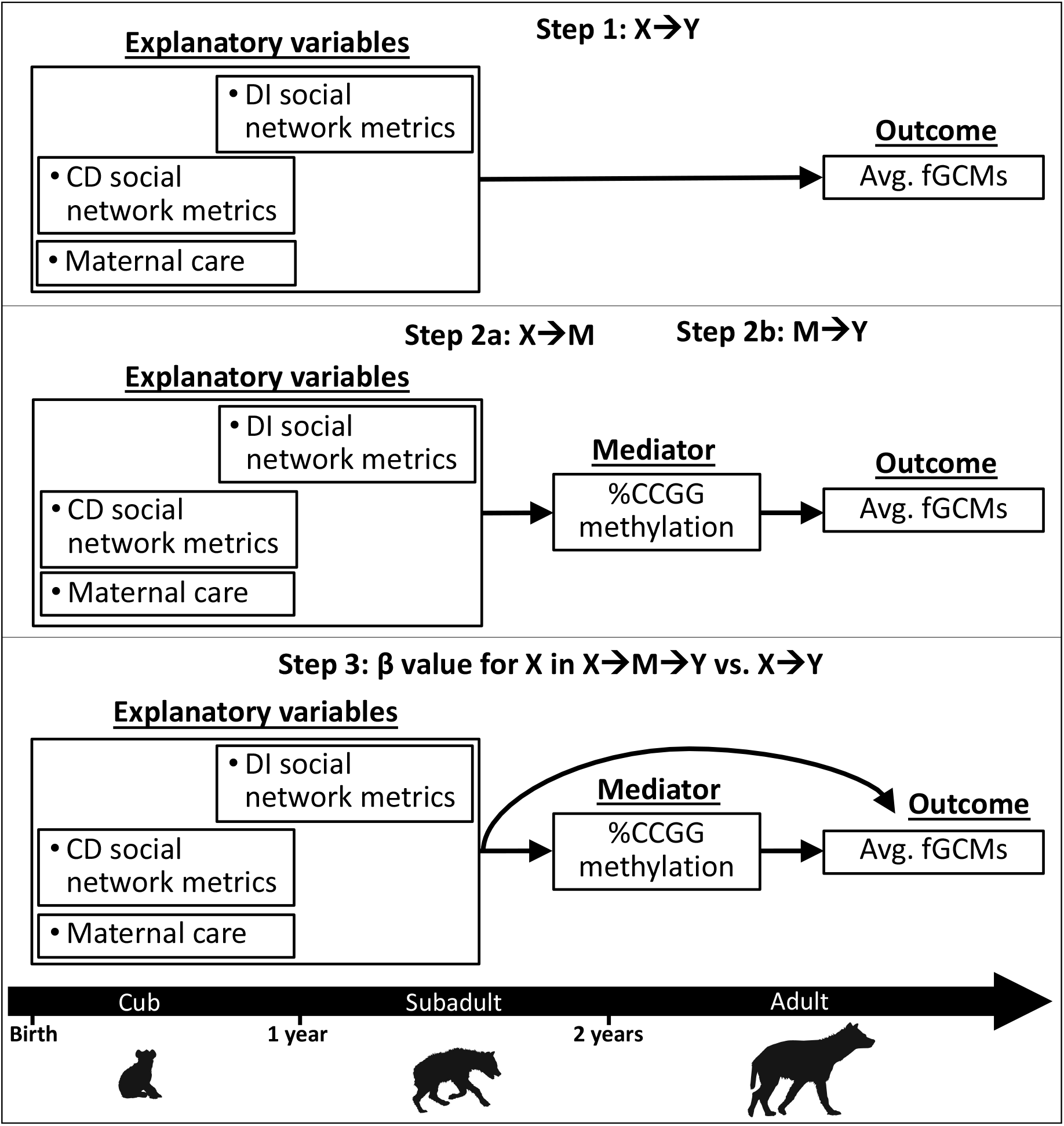
A conceptual diagram outlining the analytical strategy for the 3-step mediation analysis. X represents the explanatory variables of interest, M represents the potential mediator variable, and Y represents the outcome variable.

### Part 4: Genome-wide (mERRBS) DNA methylation analyses

Finally, we sought to identify differentially methylated CpG sites that were associated with later life corticosterone levels as well as maternal care and/or maternal social rank during a hyena’s first year of life as a way to identify novel biomarkers of this relationship. Here, we included only 30 female hyenas from a single social group born between 2011-2014 to reduce variability in background characteristics, with the exception of one hyena born in 2010 and another born in 2007. Of the selected 30 hyenas, we had information on fGCMs during the subadult or adult life stages (≥13 months) for 25 hyenas; these individuals comprised the study sample for the EWAS (**Figure 5**).

**Figure 5.**
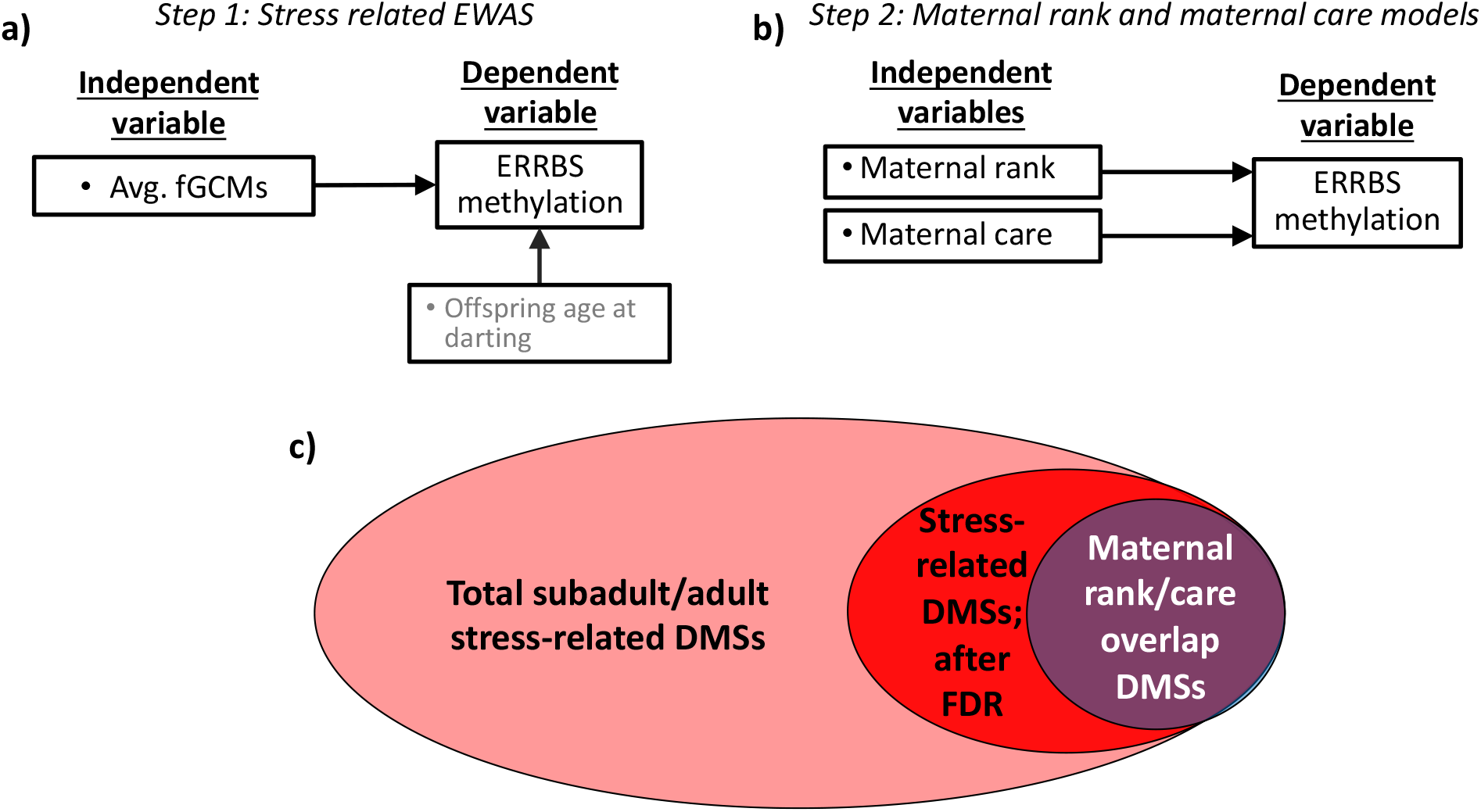
Conceptual diagram **a)** of the epigenome-wide association study (EWAS) for identification of differentially methylated sites (DMSs) from enhanced reduced representation bisulfite sequencing (mERRBS) data that are associated with subadult/adult fecal Glucocorticoid Metabolites (fGCMs) BLUPs, **b)** conceptual models of maternal rank and maternal care associations with DMSs identified in step 1, **c)** and a Venn diagram of potentially mediating stress-related DMSs that passed the false discovery rate (FDR) correction (bright red) and were also associated with maternal rank/care variables (in purple).

We carried out the final portion of the analysis following the ‘meet-in-the-middle’ approach for high-dimensional data ^38^. To prepare the fGCMs data for the EWAS, we first consolidated the repeated measures (range, 1-18) of fGCMs such that each individual had one value for stress phenotype. We used a mixed effects linear regression model where a repeated measure of fGCMs was the dependent variable, and the explanatory variable was a random effect for hyena ID and an unstructured covariance matrix. We output the best linear unbiased predictors (BLUPs) from this model, which represent the deviation of each individual’s average corticosterone concentration (i.e., the mean across all available measurements) from the population average. We then analyzed associations of these BLUPs with genome-wide DNA methylation using binomial regression models in MACAU ^39^. In these models, the BLUPs were the explanatory variable and DNA methylation of each sequenced CpG site (count of methylated divided by total CpG sites) was the outcome (**Figure 5, a**). We also controlled for offspring age in months as a covariate. In these analyses, we corrected for multiple comparisons using a Benjamini-Hochberg false discovery rate (FDR) of 5% ^40^. Among the DMSs identified in the EWAS, we further homed in on those that were also associated with maternal care and/or maternal rank. By focusing on CpG sites that are associated with later life fGCMs *and* with maternal care and/or maternal rank, we identified potential biomarkers of stress-related genomic pathways responsive to early social experience (**Figure 5, b and c**).

To accomplish this, we ran separate generalized linear regression models where the explanatory variable was individual maternal care metrics or maternal rank, and the outcomes were counts of methylated DNA sequence reads at each stress-related DMS identified from the EWAS. We treated the total number of DNA sequence read counts (methylated+unmethylated) at each DMS as an offset and assumed a Poisson distribution. We considered a CpG site to be relevant based on two criteria. First, the beta coefficient for the relationship between a given early social characteristic and DNA methylation for the CpG site of interest needed to reach statistical significance at a nominal *P*-value cut-off of <0.1. Second, given our overarching hypothesis that better quality or quantity of maternal care is protective against an adverse stress phenotype in offspring, we focused on CpG sites that exhibited opposite direction of associations for the early social metrics vs. fGCMs. That is, our hypothesis proposes that a more favorable early social environment, as indicated by maternal care, is associated with a better stress phenotype later in life, as indicated by lower fGCMs. If this is the case, a CpG site that is positively associated with maternal care (i.e., better quantity or quality of maternal care correlates with higher DNA methylation of that CpG site) should be inversely related to cortisol (i.e., higher DNA methylation of the CpG site should be associated with lower fGCMs). Thus, CpG sites of interest should theoretically be associated with maternal care and fGCMs in opposite directions in order to be potential markers of this relationship.

To interpret differentially methylated sites (DMSs) of interest (i.e., those that were identified in the EWAS, with a particular focus on those that were also associated with maternal care and/or maternal rank), we copied a 20kb sequence of hyena reference genome DNA centered on the CpG site of interest and saved the sequences as Fasta files. We first mapped each DNA sequence containing the CpG site of interest to the domestic cat (*Felis catus* Nov. 2017 [felCat9] Assembly) using the UCSC Genome Browser tool, the BLAST-like alignment tool (BLAT) ^41^, and identified nearest genes from multiple species alignment. Next we mapped each hyena DNA sequence to the human genome (UCSC Human Dec. 2010 [GRCh38/hg38] Assembly) to again identify nearest gene(s) and cross-checked the alignments with those from the more closely related cat genome.

## RESULTS

Study population characteristics and results from bivariate analyses are provided in the **Supporting Material**.

### Part 1: Associations among early social environment and adult *fGCMs*

Among hyena cubs, none of the maternal care or social connectivity measures during the communal den (CD) phase were associated with adult fGCMs levels in the adjusted models **Table 1**. However, among hyenas in the second, den independent (DI) phase of development, higher network degree (−12% [95% CI: −22%, −2%], difference in corticosterone concentration) and higher network strength (−14% [95% CI: −26%, −3%]) were associated with lower adult corticosterone levels after adjusting for a hyena’s sex, level of human disturbance in its birth year, age, reproductive state, and the time of day of sample collection **Table 1**.

**Table 1.** Association of maternal care and early life social network metrics with adult fecal Glucocorticoid Metabolites (fGCMs).

| | $\beta$ (95% CI) fGCMs<sup>†</sup> | | | |
| --- | --- | --- | --- | --- |
|  | *N* | Unadjusted model | *N* | Adjusted model<sup>‡</sup> |
| **Maternal Care FAS (per 1-SD)** |  |  |  |  |
| Close proximity | 74 | -0.13 (-0.28, 0.01) | 73 | -0.01 (-0.16, 0.13) |
| Nursing | 74 | -0.10 (-0.26, 0.06) | 73 | 0.04 (-0.11, 0.18) |
| Grooming | 74 | -0.06 (-0.21, 0.08) | 73 | 0.00 (-0.13, 0.13) |
| **CD period (per 1-SD)** |  |  |  |  |
| Degree | 79 | **-0.13 (-0.26, -0.02)** | 79 | -0.07 (-0.19, 0.04) |
| Strength | 79 | -0.11 (-0.21, 0.01) | 79 | -0.05 (-0.15, 0.06) |
| Betweenness | 79 | -0.03 (-0.19, 0.12) | 79 | -0.04 (-0.18, 0.10) |
| **DI period (per 1-SD)** |  |  |  |  |
| Degree | 79 | **-0.19 (-0.30, -0.08)** | 79 | **-0.12 (-0.22, -0.02)** |
| Strength | 79 | **-0.16 (-0.29, -0.04)** | 79 | **-0.14 (-0.26, -0.03)** |
| Betweenness | 79 | -0.10 (-0.26, 0.07) | 79 | -0.04 (-0.18, 0.11) |

<sup>†</sup> Beta estimate are fGCMs concentrations (ng/g) on the natural log scale from mixed models in which hyena ID was included as a random intercept.
<sup>‡</sup> Models have been adjusted for hyena's sex and level of human disturbance during their birth year as well as age (months), reproductive state among females, and the time of day when the fecal sample was collected.
Confidence intervals based on parametric, percentile bootstrapping; 2000 simulations.

### Part 2: Associations between early social environment and global DNA methylation

We observed multiple positive associations between early social experiences and %CCGG methylation that were robust to covariate adjustment. In our demographic covariates model, which adjusted for offspring sex and age, we found that each 1-SD greater proportion of time spent in close proximity to the mother corresponded with 1.36 (95% CI: 0.72, 2.03) greater %CCGG methylation in cub and subadult offspring. The association between close proximity and %CCGG methylation was robust to adjustment in the social experience covariates model (1.28 [95% CI: 0.34, 2.20], %CCGG), and ecological covariates model (1.07 [95% CI: 0.34, 1.83], %CCGG). In the model with mutual adjustment for all covariates, there was slight attenuation of the estimate and the confidence interval narrowly crossed the null (0.90 [95% CI: −0.05, 1.86], %CCGG, **Figure 6**). We also found that more time spent nursing was associated with higher %CCGG methylation in models that accounted for the demographic covariates (0.86 [95% CI: 0.14, 1.60], %CCGG) and social experience (1.12 [95% CI: 0.10, 2.08], %CCGG) covariates. However, adjustment for ecological covariates (human disturbance and migration status in the birth year) attenuated the effect of nursing time on %CCGG methylation. There was no effect of proportion of time spent grooming on %CCGG methylation (**Figure 6**), and maternal care and maternal rank interactions were not significant.

**Figure 6.**
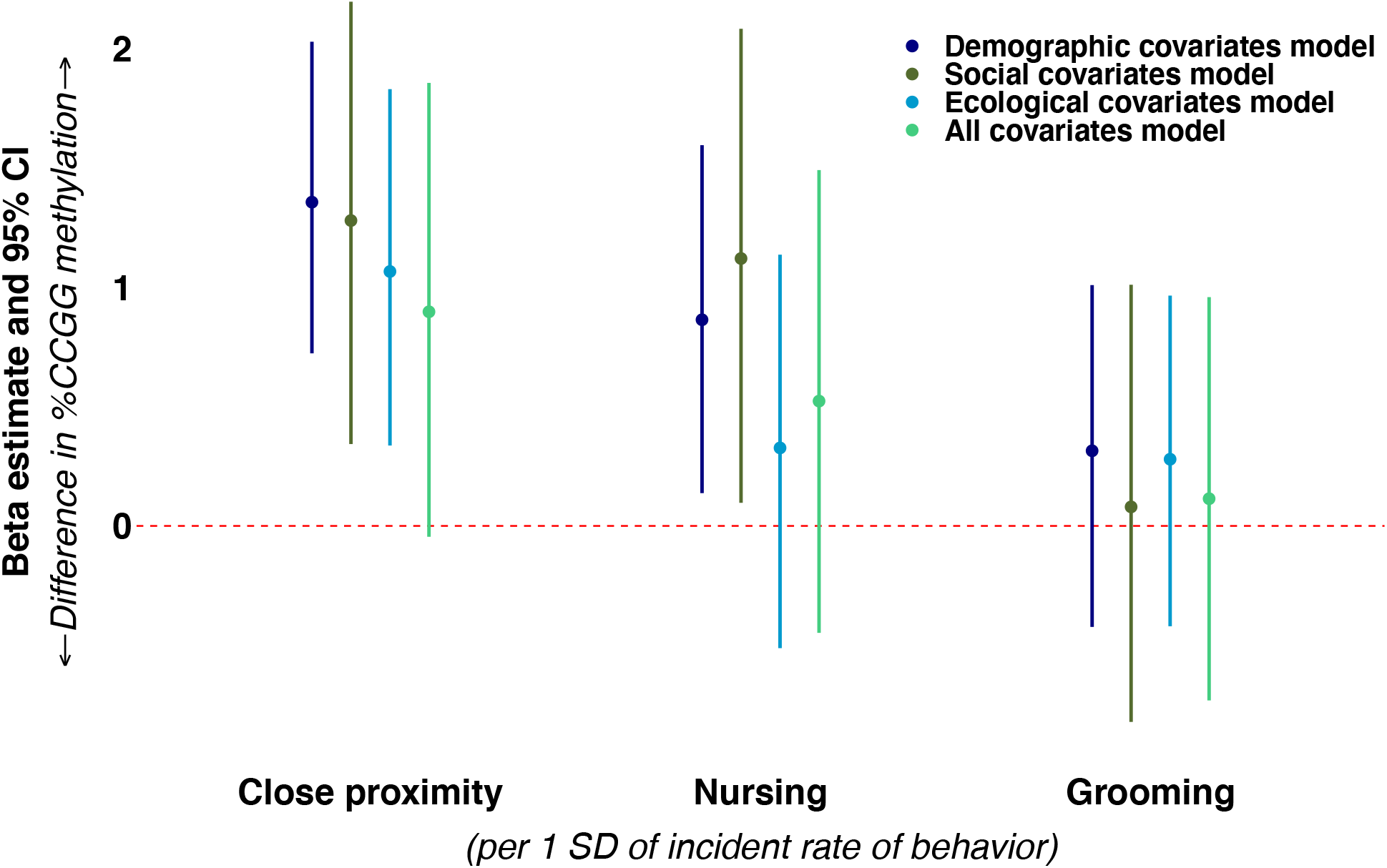
Associations of maternal care behaviors with cub and subadult offspring %CCGG methylation. All models included a random intercept for the ID of the mother and were controlled for the offspring’s age in months when the blood sample was collected for DNA methylation quantification. Models were further adjusted according to four types of covariates (demographic covariates [dark blue], early social experience covariates [dark green] and ecological covariates [light blue], and a fully adjusted all covariates model [light green]). 95% CI were based on percentile parametric bootstrapping (2000 iterations).

Next, we assessed relationships between social network metrics derived during the hyenas’ CD and DI phases of development and %CCGG. During the CD period, strength (0.75 [95% CI: 0.01, 1.48]) and betweenness (0.83 [95% CI: 0.10, 1.58]) were each positively associated with %CCGG methylation in demographic covariates models (**Figure 7**). These relationships were attenuated after adjusting for early social experience or ecological covariates. Degree was not associated with %CCGG methylation in any of our models during the CD period and none of the interaction terms involving social network metrics and maternal social rank were significant.

**Figure 7.**
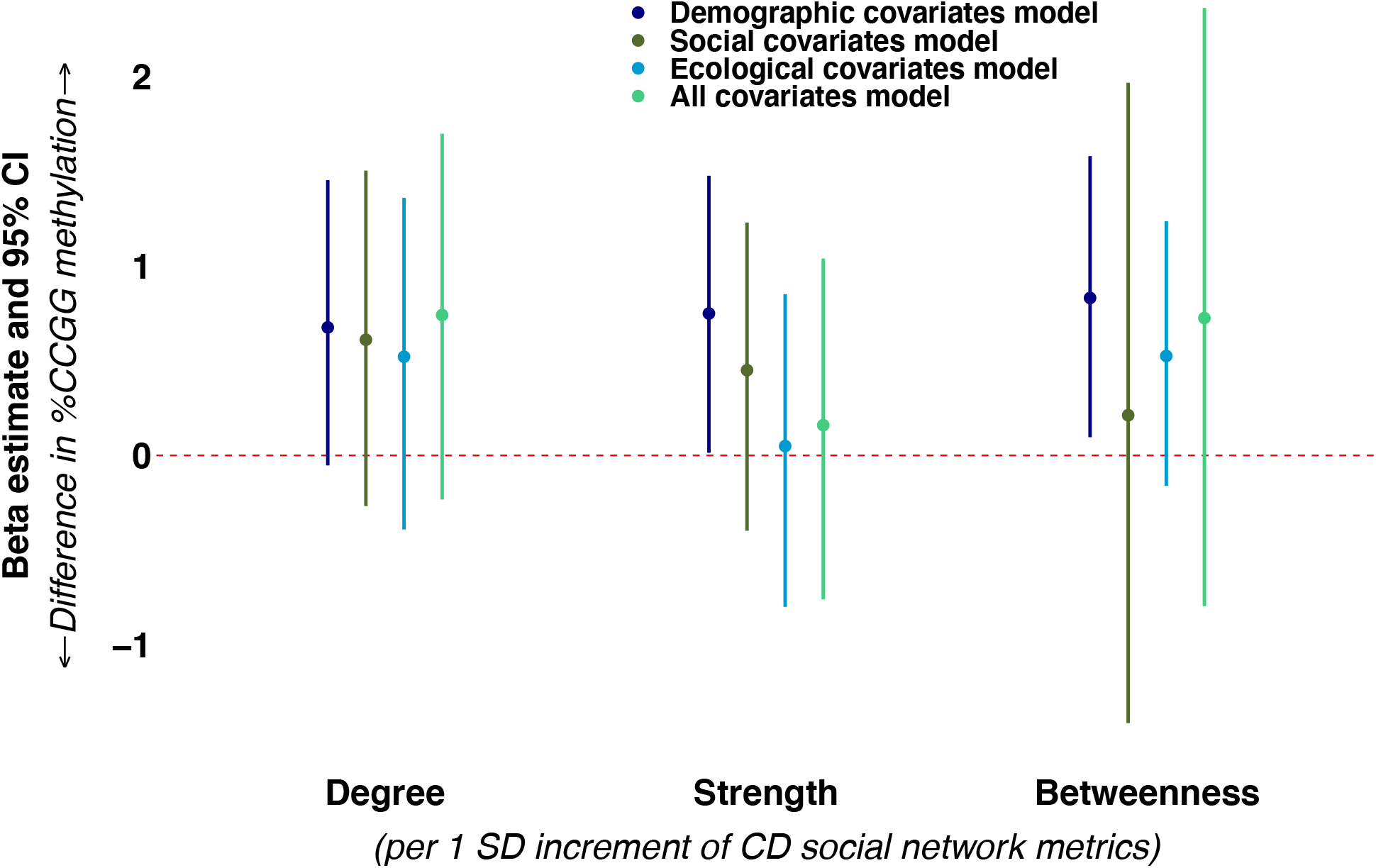
Associations of hyena CD social network metrics with cub and subadult offspring %CCGG methylation. All models included a random intercept for maternal ID and were controlled for the offspring’s age in months when the blood sample was collected for DNA methylation quantification. Models were further adjusted according to four contexts of covariates (demographic covariates [dark blue], early social experience covariates [dark green] and ecological covariates [light blue], and a fully adjusted all covariates model [light green]). 95% CI were based on percentile parametric bootstrapping (2000 iterations).

During the DI period, the degree metric was positively associated with %CCGG. Each 1-SD increment in degree corresponded with 0.69 (95% CI: 0.03, 1.38) higher %CCGG methylation in the demographic covariates model. This association was strengthened after accounting for ecological covariates (1.04 [95% CI: 0.25, 1.86] %CCGG methylation), and also in the fully adjusted model that included all covariates (1.19 [95% CI: 0.24, 2.13] %CCGG methylation). Estimates for network degree from the social experience model overlapped with the null, (0.79 [95% CI: −0.09, 1.69] %CCGG methylation, **Figure 8**.), though the direction and magnitude of estimates were similar to those of other models. Greater strength was associated with 0.71 (95% CI: 0.03, 1.43) higher %CCGG methylation in the demographic covariates model. This estimate was materially unchanged after accounting for ecological covariates (0.68 [95% CI: −0.05, 1.40] %CCGG methylation) and adjustment for all covariates simultaneously (0.83 [95% CI: −0.04, 1.72] %CCGG methylation), though we note that the 95% CIs were slightly wider in these models and narrowly crossed the null (**Figure 8**). Adjustment for social experience covariates attenuated the estimate for network strength towards the null (0.49 [95% CI: −0.33, 1.34] %CCGG methylation). Betweenness was not associated with %CCGG methylation, and there were no significant interactions between DI network metrics and maternal rank.

**Figure 8.**
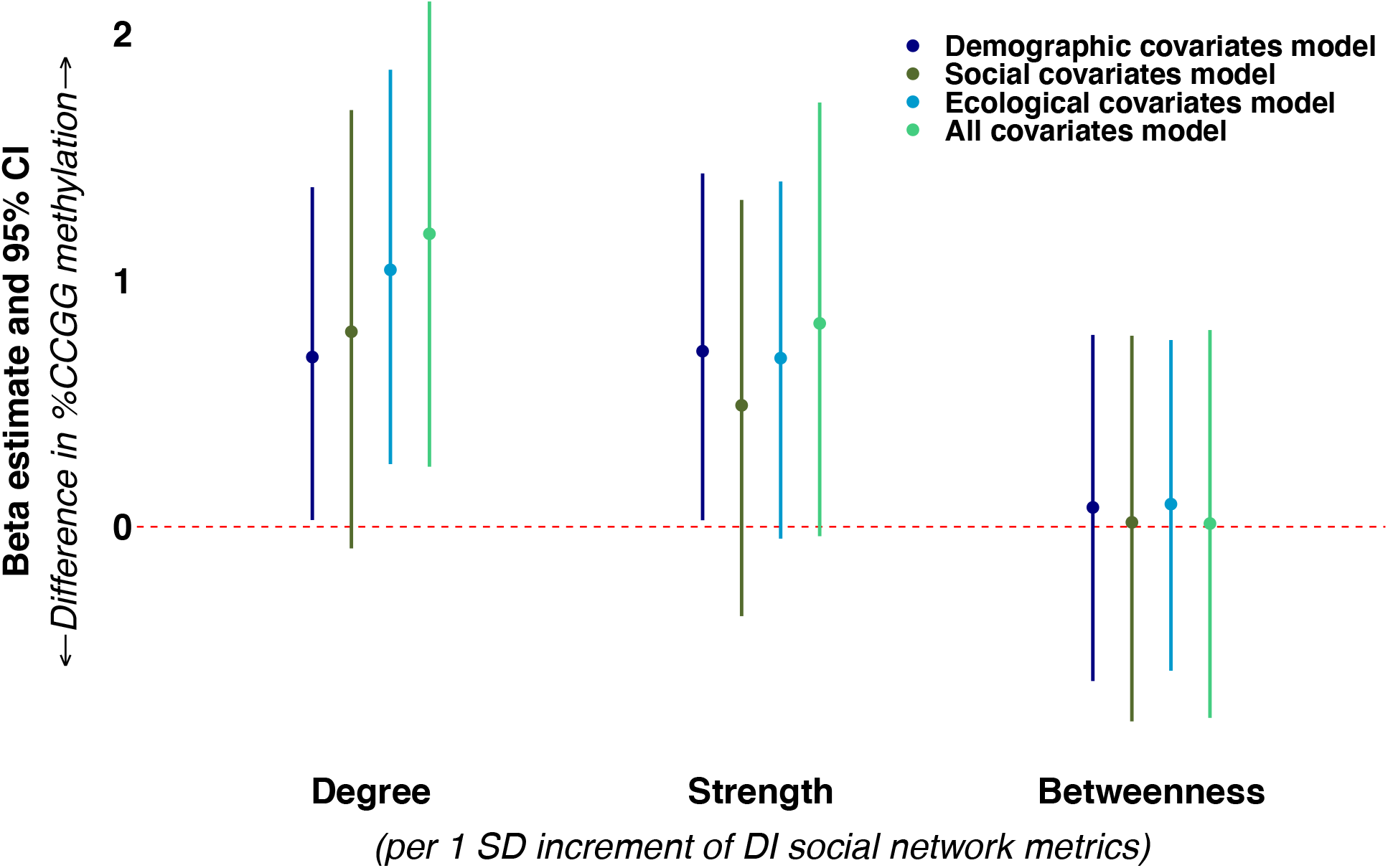
Associations of hyena DI social network metrics with cub and subadult offspring %CCGG methylation. All models included a random intercept for maternal ID and were controlled for the offspring’s age in months when the blood sample was collected for DNA methylation quantification. Models were further adjusted according to four contexts of covariates (demographic covariates [dark blue], early social experience covariates [dark green] and ecological covariates [light blue], and a fully adjusted all covariates model [light green]). 95% CI were based on percentile parametric bootstrapping (2000 iterations).

Finally, offspring %CCGG DNA methylation was not associated with adult fGCMs.

### Part 3: Formal mediation analysis

In the subset of hyenas for which we had data on maternal care (n = 30) or social network metrics (n = 52), as well as %CCGG DNA methylation and adult fGCMs, we conducted a 3-step mediation analyses. We found no association between our explanatory variables (maternal care, CD social connectedness, DI social connectedness), and the outcome, fGCMs (**Table S6**). In the second step of the mediation analyses, we found no evidence that social experiences (the explanatory variables) were associated with %CCGG methylation (the mediator), or that %CCGG methylation was associated with fGCMs (the outcome variable, **Table S6**). Given the lack of associations in the first two steps, we did not perform the final step of the mediation analysis.

### Part 4: Genome-wide (mERRBS) DNA methylation analyses

Our epigenome wide association study (EWAS) resulted in 15 DMSs that were associated with fGCMs after FDR correction (**Table 2; Figure S9**). When we examined associations between maternal care metrics and maternal rank with DNA methylation of the 15 CpG sites identified in the EWAS, five CpG sites were statistically significant. Three of these CpG sites exhibited positive *Beta* estimates for fGCMs models and corresponding negative *Beta* estimates for maternal care metrics or vice versa. At a CpG site located at the human phosphatidylinositol-4,5-bisphosphate 3-kinase catalytic subunit delta *(PIK3CD)* gene ^41,42^, the proportion of time that cubs were groomed and nursed by their mothers were each positively associated with DNA methylation (*P* = 0.007 and *P* = 0.080 for grooming and nursing, respectively; **Table 3**), and fGCMs were negatively associated with DNA methylation (*P* = 0.041; **Table 2**). We also observed a positive association of time spent grooming (*P* = 0.063; **Table 3**) and a negative association of fGCMs (*P* = 0.034; **Table 2**) with respect to DNA methylation at CpG that aligned at the lin-28 homolog B (*LIN28B*) gene in humans ^41,42^. Finally, at a CpG site that mapped between several genes, including human tubulin gamma complex associated protein 2 *(TUBGCP2)*, calcyon neuron specific vesicular protein *(CALY)*, and zinc finger protein 511 (*ZNF511)* genes ^41,42^, grooming was negatively associated with DNA methylation (*P* = 0.070; **Table 3**), and fGCMs were positively associated with DNA methylation (*P* = 0.034; **Table 2**).

**Table 2.** Epigenome wide association results showing differentially methylated CpG sites (DMSs) by fecal Glucocorticoid Metabolites (fGCMs) BLUPs among 25 female spotted hyenas. DNA methylation was assessed during the subadult life stage, and fecal samples were collected from animals during the subadult and adult life stages.

| DMSs (CpG site location<br>*C. crocuta* genome) | $\beta \pm SE$ <sup>†</sup> | FDR<br>p-value <sup>‡</sup> | Unadjusted<br>p-value | Acceptance<br>rate | Nearest human gene(s)<br>(BLAT; hg38) <sup>§</sup> | Gene position<br>(ref. hg38) | Chromosome,<br>(ref. hg38) | Human Ref. BLAT score/%<br>identical match/# bp span <sup>§</sup> | Cat Ref. BLAT score/%<br>identical match/# bp span <sup>™</sup> | Gene full name (hg38) |
| --- | --- | --- | --- | --- | --- | --- | --- | --- | --- | --- |
| C26291268 87 | -0.049 ± 0.009 | 0.041 | 2.76E-07 | 0.471 | *IQSEC1* | gene body | chr3 | 86 / 91.4% / 104 | 72 / 82.8% / 81 | IQ motif and Sec7 domain 1 |
| scaffold110 2814293 | 0.081 ± 0.015 | 0.034 | 1.18E-07 | 0.480 | *NCLN* | gene body | chr19 | 1264 / 90.9% / 21363 | 9198 / 89.1% / 19749 | nicalin |
| **scaffold13 2542930** | **-0.062 ± 0.012** | **0.041** | **2.57E-07** | **0.449** | ***PIK3CD*** | **gene body** | **chr1** | **241 / 87.4% / 996** | **9788 / 89.1% / 19872** | **phosphatidylinositol-4,5-bisphosphate 3-kinase catalytic subunit delta** |
| scaffold139 1925966 | -0.095 ± 0.018 | 0.034 | 1.21E-07 | 0.488 | *EPN2* | gene body | chr17 | 2271 / 88.7% / 25899 | 10517 / 88.1% / 21320 | epsin 2 |
| **scaffold147 1445632** | **-0.047 ± 0.009** | **0.034** | **1.46E-07** | **0.495** | ***LIN28B*** | **gene body** | **chr6** | **4774 / 87.5% / 26498** | **12257 / 91.3% / 17056** | **lin-28 homolog B** |
| scaffold193 6562606 | 0.073 ± 0.013 | 0.020 | 4.40E-08 | 0.464 | *TEX33 / TST* | overlaps both genes | chr22 | 1777 / 83.9% / 14758 | 8267 / 87.2% / 16157 | testis expressed 33 / Thiosulfate sulfurtransferase |
| scaffold207 3315565 | 0.076 ± 0.014 | 0.020 | 3.66E-08 | 0.476 | *CR1 / CD46* | between genes | chr1 | 1987 / 85.0% / 122799 | 10143 / 89.6% / 78947 | complement C3b/C4b receptor 1 / CD46 molecule |
| **scaffold248 5799761** | **0.047 ± 0.009** | **0.041** | **2.54E-07** | **0.456** | ***NDRG3 / DSN1*** | **overlaps both genes** | **chr20** | **4720 / 87.5% / 31018** | **14019 / 91.4% / 20388** | **NDRG family member 3 / Kinetochore-associated protein DSN1 homolog** |
| scaffold322 4736152 | 0.041 ± 0.008 | 0.034 | 1.64E-07 | 0.531 | *SUSD4* | gene body | chr1 | 380 / 84.4% / 8747 | 6476 / 87.4% / 16301 | single-pass type I membrane protein |
| scaffold334 1557240 | -0.080 ± 0.015 | 0.020 | 4.34E-08 | 0.434 | *MYBPC2* | gene body | chr19 | 3622 / 87.7% / 24496 | 12434 / 89.9% / 22344 | myosin binding protein C, fast type |
| **scaffold364 1284416** | **0.048 ± 0.009** | **0.029** | **7.82E-08** | **0.460** | ***WNTB2B*** | **gene body** | **chr1** | **6289 / 88.1% / 16112** | **14205 / 92.2% / 20446** | **Wnt family member 2B** |
| scaffold443 1451106 | -0.059 ± 0.011 | 0.041 | 2.45E-07 | 0.465 | *AC005258.1 / LINGO3* | gene body | chr19 | 2076 / 88.8% / 31265 | 9100 / 89.1% / 22689 | cDNA, FLJ99163 (of the GXGD family of aspartic proteases) / leucine rich repeat and Ig domain containing 3 |
| **scaffold466 8604290** | **0.033 ± 0.006** | **0.034** | **1.67E-07** | **0.478** | ***TUBGCP2 / CALY / ZNF511*** | **between genes** | **chr10** | **304 / 86.0% / 751** | **8977 / 89.0% / 41929** | **tubulin gamma complex associated protein 2 / calcyon neuron specific vesicular protein / zinc finger protein 511** |
| scaffold506 286735 | 0.068 ± 0.012 | 0.020 | 3.61E-08 | 0.437 | *DLL1* | >10kb downstream | chr6 | 927 / 88.7% / 9875 | 6679 / 87.7% / 12564 | delta like canonical Notch ligand 1 / as well as various long non-coding RNA |
| scaffold61 9436494 | -0.037 ± 0.007 | 0.020 | 3.19E-08 | 0.470 | *AC093519.2* | gene body | chr16 | 1687 / 87.4% / 23405 | 8794 / 88.2% / 20575 | methyl-CpG binding domain protein 3 |

<sup>†</sup> Untransformed (natural log scale) beta estimates are from binomial regression models run in program MACAU.
<sup>‡</sup> FDR p-value based on Benjamini-Hochberg correction (0.05%)
<sup>§</sup> Mapping of spotted hyena DNA sequences spanning 20k bp onto the human genome (Human Dec. 2013 [GRCh38/hg38] Assembly) was done using the top hit from the UCSC Genome Browser's BLAT tool.
<sup>™</sup> Mapping of spotted hyena DNA sequences spanning 20k bp onto the domestic cat genome (*Felis catus* Nov. 2017 [felCat9] Assembly) was done using the top hit from the UCSC Genome Browser's BLAT tool.
All bold DMSs were significantly associated with a maternal care metric at $P < 0.1$ .

**Table 3.**
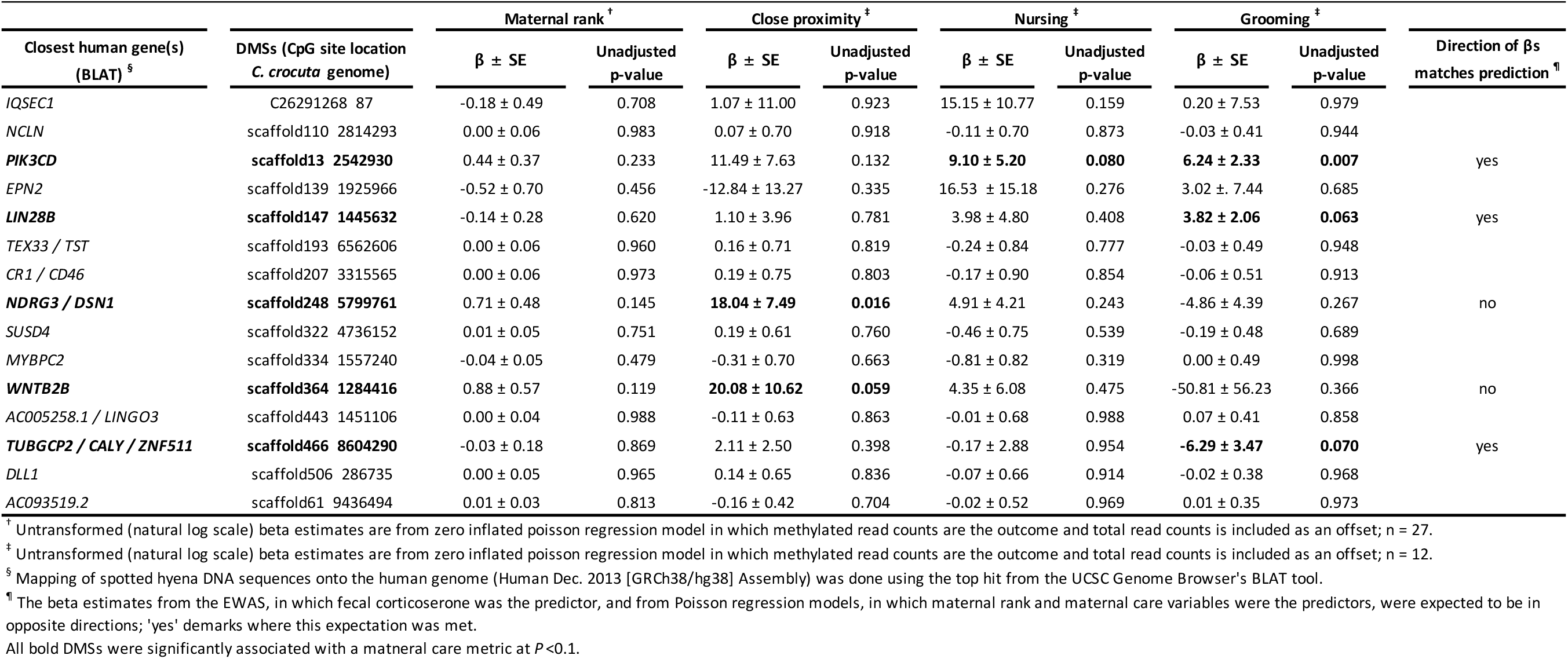
Associations of differentially methylated CpG sites (DMSs) by maternal rank and maternal care variables among the 15 DMSs identified in the stress phenotype EWAS for 12 female spotted hyenas that had no missing data (maternal rank/care, mERRBS, and fGCMs).

Additionally, two DMSs were associated with maternal care and fGCMs in which the *Beta* estimates were in the same direction. The time mother-offspring pairs spent in close proximity was positively associated with offspring DNA methylation (*P* = 0.016; **Table 3**), as well as with fGCMs (*P* = 0.041; **Table 2**) at a CpG site near the human NDRG family member 3 (*NDRG3*) and the kinetochore-associated protein DSN1 homolog (*DSN1*) genes ^41,42^. We also observed a positive association of time spent in close proximity (*P* = 0.059; **Table 3**) and a positive association of fGCMs (*P* = 0.029; **Table 2**) with DNA methylation at a CpG site that aligned near the human Wnt family member 2b (WNTB2B) gene. Despite the fact that these associations do not align with our *a priori* hypothesis that more maternal care corresponds with lower fGCMs, we believe they are worth mentioning. Maternal rank during the year in which the offspring was born was not related to any of the stress-related DMSs identified in this sample of female hyenas (**Table 3**).

### Candidate gene DNA methylation: putative glucocorticoid receptor (GR) promoter

Among 78 hyenas that passed the pyrosequencing quality checks, preliminary data revealed invariant and near zero percent methylation at six CpG sites, including the putative *NGFI-A* transcription factor binding site (**Figure S10**). Due to lack of variation in DNA methylation at these CpG sites, we did not conduct any additional analyses with respect to this region.

## DISCUSSION

Leveraging ecological and behavioral data from a long-term prospective study of wild hyenas, we investigated the extent to which social experiences during two early developmental windows are associated with DNA methylation and adult stress phenotype. We found beneficial effects on adult stress phenotype of social connectedness during the den-independent (DI) phase of development, when young hyenas no longer shelter in dens. We also found that more maternal care in the first year of life, which roughly corresponds with the CD period of development, and greater social connectedness during the DI phase were each associated with higher global DNA methylation, a presumed indicator of genomic stability and overall health. Although we found no evidence that the relationship between social experience and stress phenotype is mediated by global DNA methylation, our genome-wide DNA methylation analysis identified multiple differentially-methylated sites (DMS) that were associated with both future stress hormones and offspring receipt of maternal care, and these might serve as intermediate biomarkers for future investigation.

### The early social environment and adult stress phenotype

#### Higher social connectedness is associated with lower *fGCMs*

Greater social connectedness, specifically higher degree centrality and strength, during the DI stage when hyenas have left the communal den, corresponded with lower adult fGCMs. This supports the hypothesis that social connections promote a healthy stress response and implies that connections with group members are particularly important among subadult animals as they expand their social network beyond a limited number of associations with group members at the communal den.

Our findings corroborate previous work in humans, rodents, and baboons. In young adult humans, Kornienko et al. showed that more gregarious individuals have lower salivary cortisol levels ^43,44^. Similarly, Ponzi et al. reported that children’s perception of their social connections (i.e., higher density of friendships) were inversely related to salivary cortisol levels, as well as salivary alpha-amylase reactivity, an indicator of reactivity to psychological stressors ^45^. In Sprague Dawley rats, postweaning rearing in social isolation corresponded with higher corticosterone concentrations than in rats reared with other group members 6. The importance of social networks has also been observed in wild baboons, for which membership in more close-knit and focused grooming networks is associated with lower fGCMs ^46^. However, it is worth noting that social connections are not universally associated with stress phenotypes. A study on wild yellow-bellied marmots found no effect of social network metrics on fGCMs measured in a mixed-age, mixed-sex population ^47^.

#### Maternal care is not associated with *fGCMs*

Contrary to our predictions, and to findings from rodents, non-human primates, and humans ^3,4,7,48,49^, we found no evidence that maternal care affects future stress concentrations. There are multiple possible explanations for our null findings. First, in contrast to controlled rodent experiments, wild hyenas are subject to a multitude of stressors over development, which may hamper our ability to isolate the specific effect of maternal care on stress phenotype. Second, compared to maternal separation common in many experimental studies, natural variation in maternal care is far more subtle and may require a more sensitive measure of physiological stress than average fGCMs, which is a summary baseline stress indictor ^50^ subject to unmeasured environmental factors and the animal’s condition when the sample is collected ^51^.

### The early social environment and global DNA methylation

#### Maternal care and social connectivity are associated with global DNA methylation

Greater maternal care received during the first year of life and social connectivity during the DI life stage were both associated with higher global DNA methylation later in life. Specifically, offspring that spent more time in close proximity to their mothers during the first year of life had higher global DNA methylation later in life, even after adjustment for key demographic, social and ecological covariates. We also found that more time spent nursing corresponded to higher global DNA methylation after adjusting for demographic and social experience covariates, although the effect of nursing was attenuated when we included the early-life ecological covariates in the model. This suggests that ecological context may account for some variation in the relationship between time spent nursing and global DNA methylation, which is plausible given previously reported associations in hyenas between maternal care behavior as well global DNA methylation with respect to human disturbance and prey availability ^26,37^.

We also found that higher social connectedness during the DI, but not CD, stage of development was positively associated with global DNA methylation. Specifically, the degree of a hyena’s social connectedness, which reflects the number of group members with which a hyena associates, corresponded with higher global DNA methylation even after adjustment for demographic and ecological covariates. Controlling for early social covariates, including number of litter mates, group size, and maternal rank slightly attenuated the effect of social connections towards the null, which is not surprising given that these variables are known determinants of a young hyena’s social network ^21^ as well as of global DNA methylation ^26^.

### Mediation by global DNA methylation

#### No evidence of mediation by global DNA methylation

We found no evidence of mediation by %CCGG methylation. This could be due to the fact that %CCGG methylation, averaged over the entire genome, may be too broad a metric to isolate important regulatory pathways involved in the stress response; it might also be due to low power given our relatively small sample sizes in the mediation analysis. Therefore, we complemented this analysis with Reduced Representation Bisulfite Sequencing (mERRBS) ^52^, a high throughput sequencing technology that assays single nucleotide DNA methylation resolution at a genome wide scale.

#### Biological interpretation of CpG sites identified in genome-wide DNA methylation assessment

Using mERRBS, we identified 15 DMSs that were associated with fGCMs after FDR correction, five of which were also associated with early life maternal care metrics. Three of these DMS followed our prediction given that effective maternal care is hypothesized to be protective against an adverse stress phenotype. More specifically, for a given DMS, a positive association between DNA methylation and stress hormone levels should correspond to a negative association between DNA methylation and maternal care at that DMS, or vice versa. We discuss the direction of the associations of these three DMSs with respect to fGCMs and early life maternal care metrics below, as well as their potential biological function.

We found that offspring DNA methylation of a CpG site that mapped near the human *PIK3CD* gene was both positively associated with time spent nursing and receipt of maternal grooming, and inversely associated with fGCMs. This gene is highly expressed in leukocytes where its lipid product is a critical signaling component of immune response ^53,54^. More specifically, *PIK3CD* is involved in the development of immune cells, particularly B cells as well as natural killer cells ^55,56^. Pathophysiology resulting from under and over expression of this gene range from immune deficiency disorders to autoimmune disorders and cancers ^57^. Second, we identified a CpG site that mapped near the human *LIN28B* gene and was positively associated with receipt of maternal grooming and inversely associated with fGCMs. In a rat model of depression, animals with depressive phenotypes exhibited higher expression of *LIN28B* and *IL6* mRNA. This suggests that LIN28B is involved in the *IL6* regulatory pathway ^58^, which has been implicated in chronic stress and depression pathways in both animals and humans ^59,60^. Additionally, two recent meta-analyses of genome-wide association studies linked differential expression of *LIN28B* to depression in multiple large human cohorts ^61,62^. Finally, genetic variation of the *LIN28B* gene is associated with age at menarche in humans ^63^. Early social stressors have also been found to be associated with age of menarche ^64^, which in turn was associated with depressive symptoms ^65,66^, suggesting that a pathway involving the *LIN28B* gene may link early social environment, growth and development, and stress phenotypes.

Lastly, we found that DNA methylation of a CpG site between human *TUBGCP2, CALY, and ZNF511* genes was inversely associated with receipt of maternal grooming and positively associated with fGCMs. Given the uncertainty about the location of this CpG site, we reserve speculating about its potential biological function. Taken together, despite having only correlative evidence from our EWAS, the identification of a potential molecular biomarker in hyenas near the human *PIK3CD* and *LIN28B* gene suggests that pathways involving immune function and inflammation may be important for understanding the relationship between early life maternal care and later life stress phenotypes.

### No evidence of DNA methylation variation at the putative hyena glucocorticoid receptor (GR)

Although some previous work in rodents and humans has shown that inadequate maternal care and greater early adversity corresponded with higher methylation of CpG sites the GR promoter region ^10,67^, the inverse relationship between maternal care and GR DNA methylation has not been replicated in all studies ^68^. In the present study, we measured DNA methylation at CpG sites in the putative hyena *GR* gene promoter and found invariant and near zero percent methylation, suggesting a potential lack of plasticity in DNA methylation in this promoter region, at least when using peripheral leukocytes. Future studies may better replicate results from laboratory rodents if using brain tissues where expression of the *GR* gene is more tightly coupled with HPA regulation.

## CONCLUSIONS

In this wild population of spotted hyenas, maternal care and social connections early in life were key determinants of global DNA methylation and future fGCMs, even after accounting for key demographic, biological, and social factors. These findings contribute to the literature in multiple ways. First, few studies have been able to assess early social experience, DNA methylation and future stress phenotype together, a study design that allows for explicit assessment of DNA methylation as a potential mediator. Second, existing literature has focused predominantly on single social experiences during specific developmental windows, which is not only unrealistic given the multitude of experiences across the life course that may contribute to stress phenotypes, but also precludes the ability to assess the relative importance of the type and timing of experiences. In the present study, we were able to examine prospective associations of multiple aspects of social experience during two vulnerable windows of development. Finally, to our knowledge, this is the first study in a wild animal system that has considered the proximate role of DNA methylation as a mechanism that may facilitate developmental plasticity of a stress phenotype in response to maternal care and social connections with group members.

Nevertheless, our study has limitations. We used a large database of existing behavioral, demographic and biological data to test our hypotheses. As a result, we lacked complete data overlap between different variables of interest, and thus models in different parts of our analyses are not directly comparable. We also took the average of multiple measures of fGCMs as an indicator of the animals’ stress phenotype. However, a more nuanced measure of stress, like reaction and recovery to a standardized stressor, may provide a more sensitive assessment of acute HPA function ^69^, while fGCMs capture average corticosterone levels over time which may be more relevant to chronic stress-related phenotypes. Finally, our mediation and mERRBS analyses comprised small sample sizes, and therefore may have low external validity – particularly for the mERRBS analysis, which included only females. Future analyses would benefit from a well-annotated hyena genome and the capacity to filter for SNPs, information on cell type heterogeneity, and inclusion of a pedigree to control for genetic correlations in DNA methylation.

Our work with hyenas joins a growing body of evidence from rodents and primates highlighting the critical effects of early experience on molecular biomarkers and future stress phenotype. Given that such studies are seldom possible in free-living populations of long-lived, gregarious animals, this study provides novel insight on the type, timing, and mechanisms that link early-life experiences with the developmental plasticity of stress phenotypes and highlights the extent to which natural variation in maternal care and social interactions have persistent effects on developing offspring.

## ACKNOWLEDGEMENTS

We thank the Kenyan National Commission for Science, Technology and Innovation, the Kenya Wildlife Service, the Narok County Government, and the Senior Warden of the Masai Mara National Reserve for permission to conduct this research. We are indebted to all those who have contributed to long-term data and sample collection on the Mara Hyena Project. We are also grateful to Dr. Elise Zipkin for suggestions and feedback regarding the statistical modeling, as well as Dr. Lu Tang and Patrick Bills for help processing the behavioral data. This work was supported by National Science Foundation Grants DEB1353110, OISE1556407, and IOS1755089 to KEH, and Doctoral Dissertation Improvement Grant from NSF (DDIG 1701384) to ZML. This work was also supported in part by funds from NSF Grant OIA 0939454 to the BEACON Center for the Study of Evolution in Action as well as Michigan Lifestage Environmental Exposures and Disease (M-LEEaD), NIEHS Core Center (P30 ES017885), as well as the UM NIEHS Institutional Training Grant T32 ES007062 to DCD.

