## Supporting Material for "Associations of early social experience with offspring DNA methylation and later life stress phenotype"

**Supporting Information**

**METHODS**

*Maternal care behaviors from focal animal survey (FAS)*

Our maternal care data included 1533 FAS totaling approximately 779 hours of observations of 258 mother-infant pairs when offspring were 1 year old or younger, which is the approximate age at weaning [1]. We also filtered these data to include only FAS sessions during which the mother was lactating, and the mother and offspring were observed together for a minimum of five minutes. Given that we had anywhere from 1 to 29 repeated observations of mother-offspring interactions for each maternal behavior of interest (close proximity, nursing, and grooming) for each mother-offspring pair, the first step to processing these data for analysis was to convert the repeated measurements into a single value for use in the regression models. To do this, we fit generalized linear mixed-models where the repeated outcome of interest was counts of each maternal care behavior for a given mother-offspring pair. Fixed effects covariates included key characteristics having potential to affect mother-offspring interactions: the offspring's age in months on the date of the FAS, timing of the FAS (morning or evening), and FAS season (during the annual wildebeest and zebra migration, present or absent). We also included an offset of the natural log of the length of time the mother-offspring were both present during the FAS to control for observer effort. (*Nota bene*: the offset also facilitates interpretation of parameter estimates such that they reflect incident rates or proportions of time spent engaged in a particular behavior in relation to the total time the mother-offspring pair was observed). Our models also included a random intercept for offspring hyena ID to account for correlations among the repeated observations. Using these mixed models, we calculated the Best Linear Unbiased Predictors (BLUPs) for each mother-offspring pair. Because behavioral count data are often over-dispersed and zero-inflated, we fit models assuming three different underlying distributions each with and without a zero-inflation correction for a total of six model specifications per type of maternal care behavior. We used the R package glmmTMB to fit a Poisson distributed model in which the mean equals the variance, as well as two parameterizations of negative binomial distributed models which differ in how the variance scales with respect to the mean [2,3]. More specifically, in the negative binomial 1 models, the variance is scaled as multiplicative function of the mean and an estimated dispersion parameter, while in the negative binomial 2 models the variance is scaled as a quadratic function of the mean and an estimated dispersion parameter [4]. We used a simple form of a zero-inflation correction in which each observation has an equal probability of being a zero [2]. We compared model fit using AIC for each distribution with and without modeling zero-inflation. After selecting the best-fitting model with the lowest AIC (**Supporting Table 1**), we extracted the individual-level random effect estimates from models in which close proximity, nursing, and grooming were the outcomes. These individual-level random effect estimates represent the underlying distribution of deviation of each maternal care behavior received by each hyena in comparison to the population average for that particular behavior. We then added the individual random effects to the overall model intercept, which represents a relevant population-level biological anchor (i.e., the average amount of maternal care received by the overall population when all other variables are set to the referent level). After appending the BLUPs to the intercept, we exponentiated the variable to transform the estimates from the natural log scale back to the original scale, proportion of minutes during which the mother-infant pair were observed together. Finally, we z-score standardized all maternal care BLUPs prior to using these measures as explanatory variables in downstream models.

**Supporting Table 1.** AIC values for generalized linear mixed models and zero-inflated generalized linear mixed-models of maternal care behaviors. The best fitting model (lowest AIC) was used to generate Best Unbiased Linear Predictors (BLUPs) for each hyena offspring.

| Maternal Care behaviors | dAIC | Deg. freedom |
|---|---|---|
| **Close proximity** | | |
| Zero-inflated negative binomial (1) distributed model †,‡ | 0.0 | 7 |
| Zero-inflated negative binomial (2) distributed model †,§ | 3.3 | 7 |
| Negative binomial (1) distributed model | 468.3 | 6 |
| Negative binomial (2) distributed model | 611.4 | 6 |
| Zero-inflated Poisson distributed model † | 1563.1 | 6 |
| Poisson distributed model | 4601.4 | 5 |
| **Nursing** | | |
| Zero-inflated negative binomial (1) distributed model †,‡ | 0.0 | 7 |
| Zero-inflated negative binomial (2) distributed model †,§ | 17.0 | 7 |
| Negative binomial (1) distributed model | 773.4 | 6 |
| Zero-inflated Poisson distributed model | 791.6 | 6 |
| Negative binomial (2) distributed model | 1062.9 | 6 |
| Poisson distributed model | 9518.6 | 5 |
| **Grooming** | | |
| Zero-inflated negative binomial (1) distributed model †,‡ | 0.0 | 7 |
| Negative binomial (1) distributed model | 16.7 | 6 |
| Zero-inflated negative binomial (2) distributed model †,§ | 32.2 | 7 |
| Negative binomial (2) distributed model | 39.2 | 6 |
| Zero-inflated Poisson distributed model † | 113.7 | 6 |
| Poisson distributed model | 739.4 | 5 |

Models are adjusted for offspring hyena's age (months), time of day, and migration season on the date of the FAS.
Models include a random intercept for offspring ID.
† Single zero inflation parameter applied to all observations.
‡ Negative binomial 1 where, variance = $(1+\alpha)\mu$.
§ Negative binomial 2 where, (variance = $\mu + \alpha\mu^2$

Our FAS data were collected during four distinct study periods between 1998 and 2013. Therefore, we checked for consistency between the four periods of data collection by visually inspecting a principle component analysis (PCA) plot (**Supporting Figure 1**). We saw no evidence of clustering by sample collection period and concluded that variation in maternal care was likely not confounded by sampling batch effects, so we were able to pool data from all study periods together.

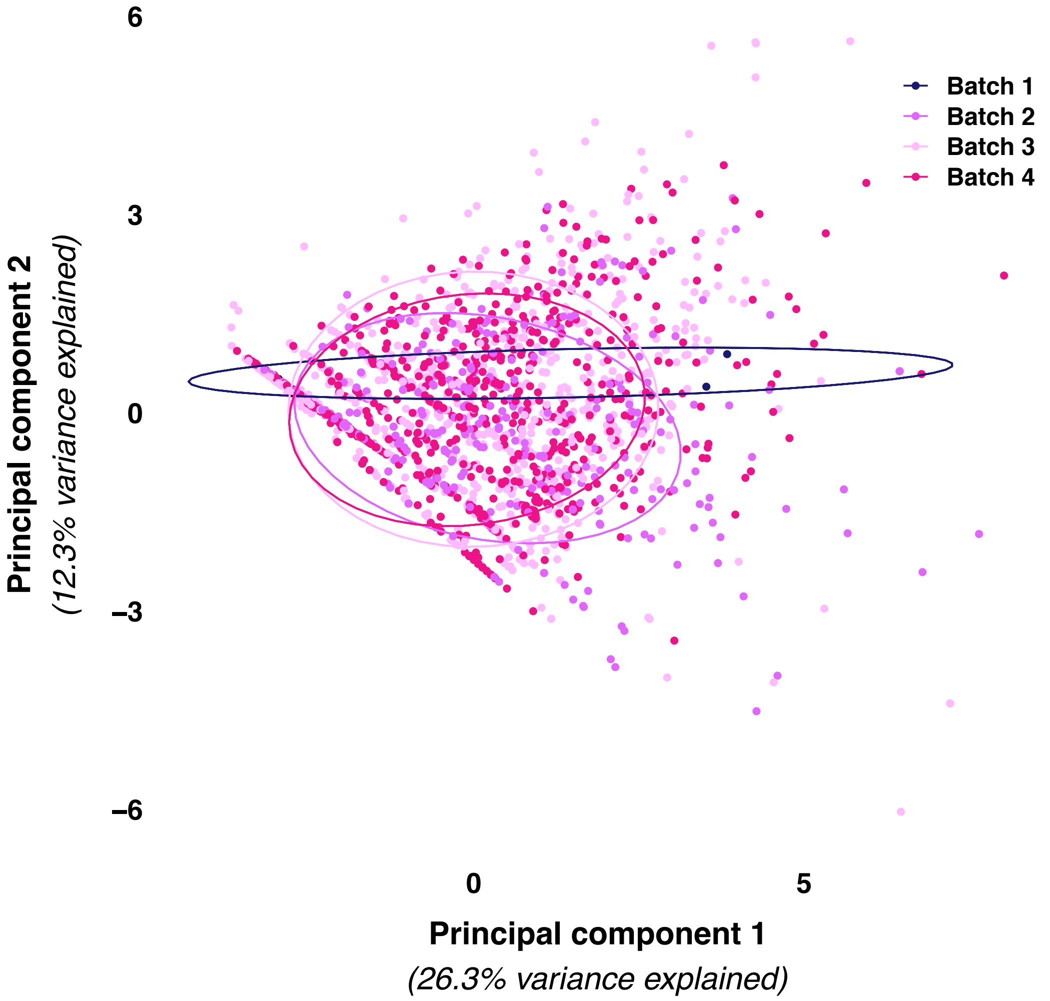


**Supporting Figure 1.** Principal Components Analysis (PCA) plot of focal animal survey (FAS) batches (data collection groupings) for maternal care behaviors. Ellipses represent 68% probability. Batches represent unique study periods. A square root transformation was used on count data prior to the PCA.

*Social network metrics derived from 115 hyenas during the communal den dependent (CD) and communal den independent (DI) phases of development*

Association social networks were based on regular 15-20 minute interval scan sampling during observations sessions in which two or more hyenas were present [5]. The networks incorporated twice weighted association index data [6], which enabled us to correct for sampling bias that could stem from variation in hyena group observability, thus providing more reliable information about social bonds in our population [5,7]. In our association networks, degree centrality (i.e., degree) corresponded to the number of different individuals with which the hyena was recorded in the same session. Strength, a weighted metric of degree or network connectedness [8], corresponded to the total number of times the hyena was observed with other clan mates, including repeated associations with the same individuals. Betweenness centrality (i.e., betweenness) was calculated as the number of shortest paths that connected hyenas in the group and passed through the hyena of interest [8]. An individual with high betweenness can be thought of as a bridge in which otherwise unconnected members of the network are indirectly connected. As with maternal care, we z-score standardized all association index social network metrics prior to using these measures as explanatory variables in downstream models.

*Blood collection, processing, and storage*

Hyenas from our study population were immobilized using a $CO_2$ rifle that propelled a pressurized dart containing 6.5 mg/kg of tiletamine-zolazepam (Telazol ®). Within 13 minutes of when the hyena was sedated, we drew blood from their jugular vein into ethylenediaminetetraacetic acid (EDTA) coated vacuum tubes. We flash froze whole blood samples in liquid nitrogen or we extracted genomic DNA using Gentra Pure Gene or PAXgene Blood DNA kits by Qiagen® and stored samples until they were transported to the U.S. for long term storage at -80°C.

*Global (%CCGG) DNA methylation*

We used the LUminometric Methylation Assay (LUMA) [9,10] to measure global DNA methylation. Our laboratory procedure, and data processing have been previously described [11]. In brief, this assay uses a parallel enzyme digestion of DNA to quantify the amount of methylated vs unmethylated CpG sites within the '5-CCGG-3' recognition sequence. Following pyrosequencing of the enzyme-digested DNA, we calculated a composite global methylation value for each hyena representing the average DNA methylation across the hyena genome. As we and others have previously described, there are approximately 2.4 million CCGG motifs in the mammalian genome, and based upon the high resolution human genome, we expected about 3% of the CCGG motifs to occur within 1kb of transcription start sites, 45% in gene bodies and 52% in non-coding regions of the genome [11–13].

*Genome-wide DNA methylation: Enhanced Reduced Representation Bisulfite Sequencing (mERRBS)*

Genomic library preparation and sequencing

First, we digested approximately 100ng of genomic DNA for 16-18 hours with the restriction enzyme MSpI. Digested DNA was purified via phenol-chloroform extraction and ethanol precipitation. Next, we ran the digested DNA through an end-repair reaction, and an A-tailing reaction, which adenylated the 3’ of the DNA. Third, we ligated paired-end methylated adapters to the A-tailed DNA and incubated at 16°C overnight [14]. Next, we selected the ligated DNA fragments in the ranges of 150-250 bp and 250-450 bp from an agarose gel and purified the excised fragments with a Qiagen QIAquick® Gel Extraction kit. Size-selected DNA was bisulfite treated using Zymo Easy DNA Methylation™ kits, followed by PCR enrichment of the bisulfite converted DNA using the Roche FastStart™ High Fidelity PCR system. Prior to sequencing, DNA libraries were cleaned using AMPure XPSPRI beads, and DNA library quantity and quality were assessed with Qubit® High Sensitivity dsDNA kit and Agilent’s High Sensitivity D1000 Tape screen, respectively. Finally, we multiplexed five libraries per flow cell on an Illumina HiSeq4000® platform for single-end sequencing with a 50-nucleotide read length. Included alongside of the hyena samples were libraries for a human genomic DNA sample and hyena samples were spiked with a lambda phage DNA sample to serve as a control and to estimate bisulfite conversion efficiency. More detailed information on the mERRBS library preparation and sequencing protocol, can be found in Garrett-Bakelman et al. (2015).

Bioinformatics pipeline

We assessed the quality of raw mERRBS data and identified specific reads from each sequenced sample that required trimming using FastQC (FastQC) (v0.11.3). We used TrimGAlore (Trim Galore!) (v0.4.5) to remove low quality bases with Phred quality scores < 20, as well as adapter sequences, primers and extra bases from the 3’ ends of reads that were the product of poly-A tail end-repair. Next, we used Bismark (v0.19.0) to perform alignment and methylation calling (identification of methylated vs unmethylated CpG sites) of sequenced short reads [16]. In this program, residual cytosines are converted to thymines in both the sequenced reads and the reference spotted hyena genome provided by the Bejing Genome Institute. Using Bowtie2 (v2.3.4) [17] within Bismark, we then aligned short reads using the default parameters (multi-seed length of 20bp with 0 mismatches) and methylation calls were retained for all nucleotides with a read depth ≥ 5. Prior to downstream statistical analysis, we further restricted the data set to nucleotides with a read depth ≥ 10 and we used methylKit to exclude nucleotides that did not have coverage across all samples [18].

*Candidate gene DNA methylation*

We used a target gene approach to measure DNA methylation of CpGs within the putative promoter region of the spotted hyena glucocorticoid receptor (GR) gene. To do this we *de novo* sequenced the GR promoter region of DNA in hyenas. Given that there was no publicly available hyena genome at the time, we first made an in silico 'synthetic strand of DNA,' which was intended to represent our best guess at the nucleotide sequence for hyenas and which was based on the consensus base pairs from the alignment of multiple species' GR promoter sequences (**Supporting Figure 2**). More specifically, our multiple species alignment contained human (including from McGowan *et al.* 2009), domestic dog, and walrus GR promoter DNA. Then moving across the alignment, we selected the nucleotide for our 'synthetic strand of DNA' at each position based on the greatest nucleotide similarity across species or based on the nucleotide from species most closely related to hyenas when there was no clear consensus.

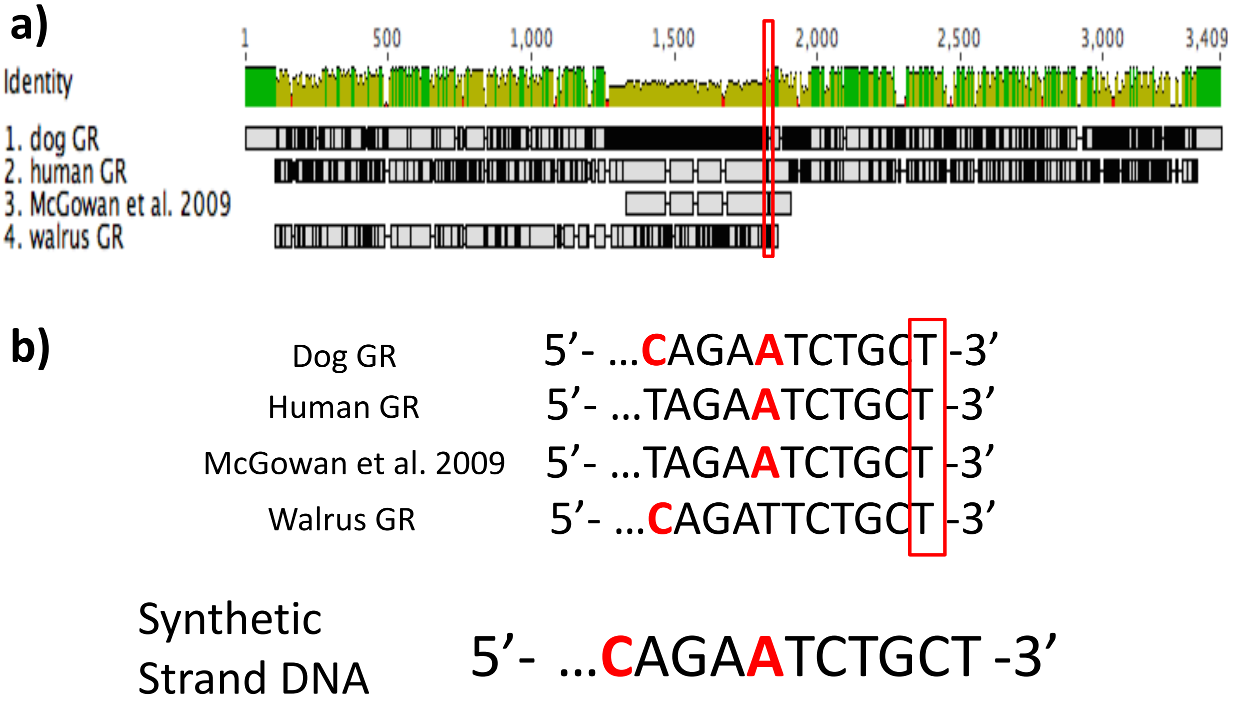


**Supporting Figure 2**. (a) A multiple species alignment of the promoter region of the Glucocorticoid Receptor, where consensus nucleotide identity is represented by the green bar at the top. (b) A zoomed in view of the nucleotides from each species in the alignment and the 'synthetic strand of DNA' at the bottom. Red boxes represent the sliding window used to assess nucleotide consensus at each position and red nucleotides indicate a mismatch between one or more species.

Next, using our synthetic strand of DNA, we designed a tiling array of PCR primers with overlapping amplification products that targeted the GR promoter region, including parts of this promoter that had been shown to be differentially methylated in human and rodent studies (c.f. [19,20], **Supporting Figure 3**). We size selected spotted hyena DNA PCR products from gels based on the estimated amplicon lengths from our 'synthetic strand of DNA,' and Sanger sequenced the size matched products (**Supporting Figure 4**). In Sanger sequencing, we used genomic DNA extracted from whole blood samples taken from three individual hyenas.

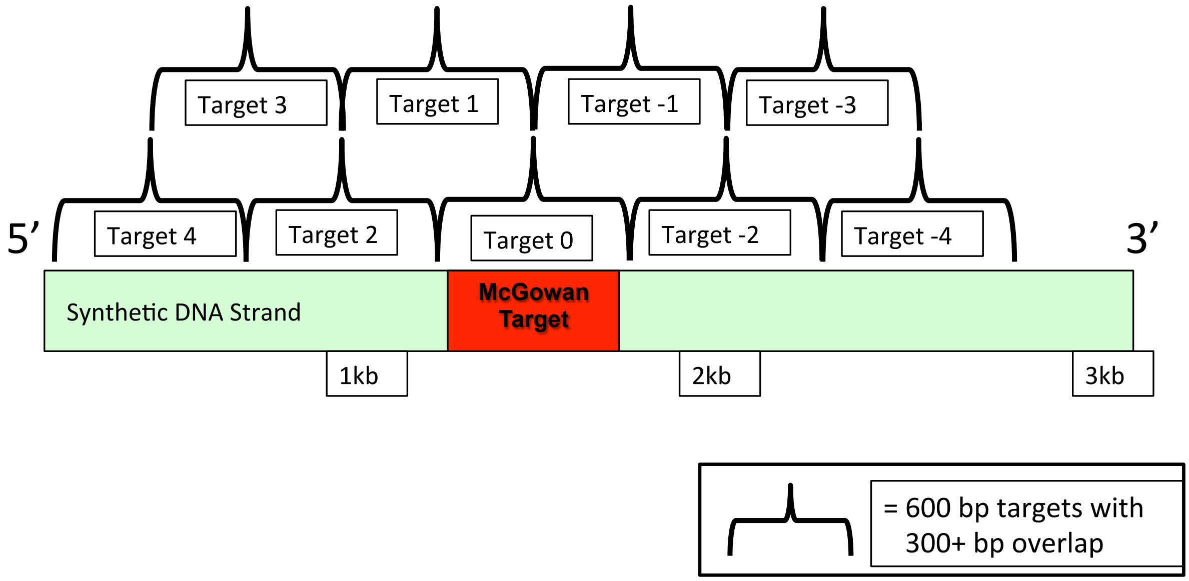


**Supporting Figure 3**. Tiling array of PCR primers and targets aimed at generating overlapping DNA sequence from spotted hyenas that aligned with GR promoter region and included sequence of interest identified in human DNA methylation studies (e.g. McGowan target).

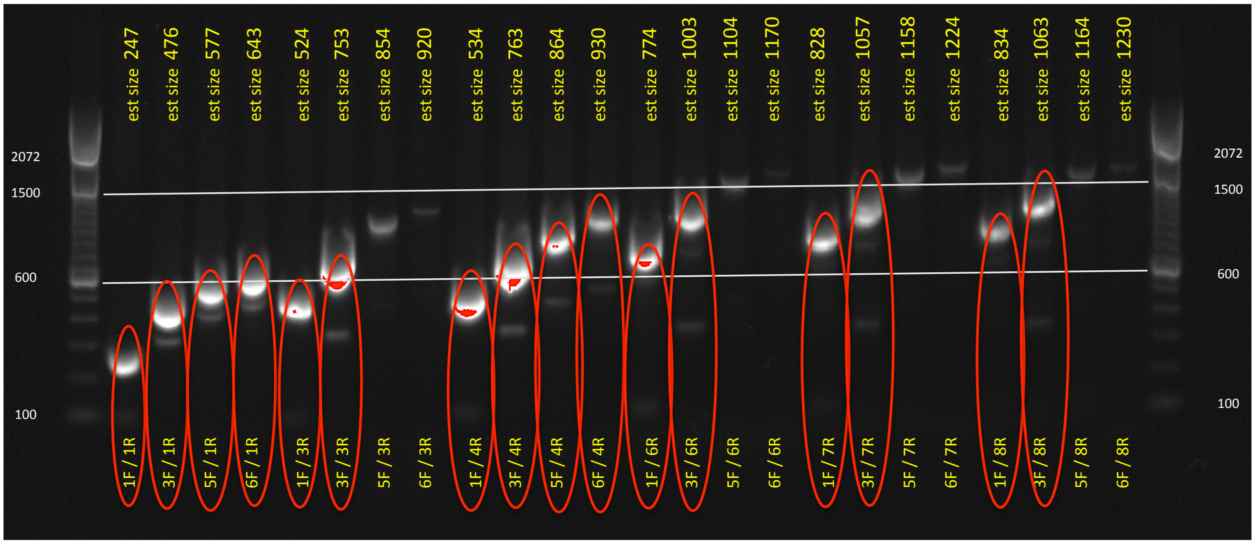


**Supporting Figure 4**. A gel image showing the PCR products from our tiling array. Size ladders are shown in the first and last wells. Primer sets, identified on the bottom of the gel, and their respective DNA bands are circled in red if the PCR product matched the estimated size. Estimated sizes were based on the ‘synthetic strand of DNA’ and are listed across the top of the gel image.

We then aligned the overlapping hyena DNA reads from the Sanger sequencing and trimmed the hyena DNA to match published human GR DNA sequence [20–22], which resulted in the assembly of the putative GR promoter region for spotted hyenas (**Supporting Figure 5**).

#### Putative spotted hyena GR Promoter

**5' -** gggttctgctttgcaacttctctcccggtgggagagcgcgacggcggcgg
cggctgcagacgggg**cCGccc**agacgatgcggcggtgggggacctgccgg
cacgcgactacccccgagtcgagagtatgtacgcaccgacccccctcctct
ctcctcccctccctcagcctccccagagggcgtgtctggctgtccggccc
cgagcgaggccgagacgctgcggcaccgtttccttgcaaccctcgtagcc
ccgtgctaagtgacacagttcgcgcaactccgctccgaggcggcggcgcg
gaccactcccgcggccccgctgcgggccgcgccccggcctccccccaccc
cccacc**cCGccc**cagacagcccgctgtcacccgcaggggcgctgacggcg
ccggtcgccgagggactgggccagctccggagtgggtcgggagtcgcgcc
gctgggctg**gggCGg**g*aaggaggcagcgaggagagaaactagagaaactc*
*ggcttccctcccaagctcg*ccccggagagaccaggtcggcctccagccgc
cacctctccctttcccggagggtgggggg **- 3'**

**Supporting Figure 5**. Spotted hyena putative GR promoter sequence, which has been trimmed to match the aligned sequences published in Oberlander et al. 2008, McGowan et al. 2009, and Perroud et al. 2011. Underlined is the inferred consensus region (based on alignment) for human exon $1_F$, and red, italicized is the inferred consensus region (based on alignment) for rat exon $1_7$. GC boxes, the putative binding site for transcription factor NGFI-A, are **bold** and contained CpG sites capitalized. The yellow highlighted GC box is the potential NGFI-A transcription binding site homologous to the region in rats that was shown to be differentially methylated with respect to maternal licking and grooming (c.f. Weaver et al. 2004).

After sequencing the putative GR promoter in hyenas, we aligned our trimmed sequence with human and rat DNA sequences that were reported in the literature to be differentially methylated with respect to early life adversity and maternal care, respectively [19–24] and as shown in **Supporting Figure 6**. Focusing on an approximately 50 base pair region which overlapped with the differentially methylated regions in human and rat studies, including the putative transcription factor binding site for NGFI-A, we quantified CpG methylation using pyrosequencing on a Qiagen Pyromark® Q96 MD. We bisulfite treated genomic DNA extracted from whole blood, and then measured % DNA methylation in 96 hyena samples at 6 CpG sites.

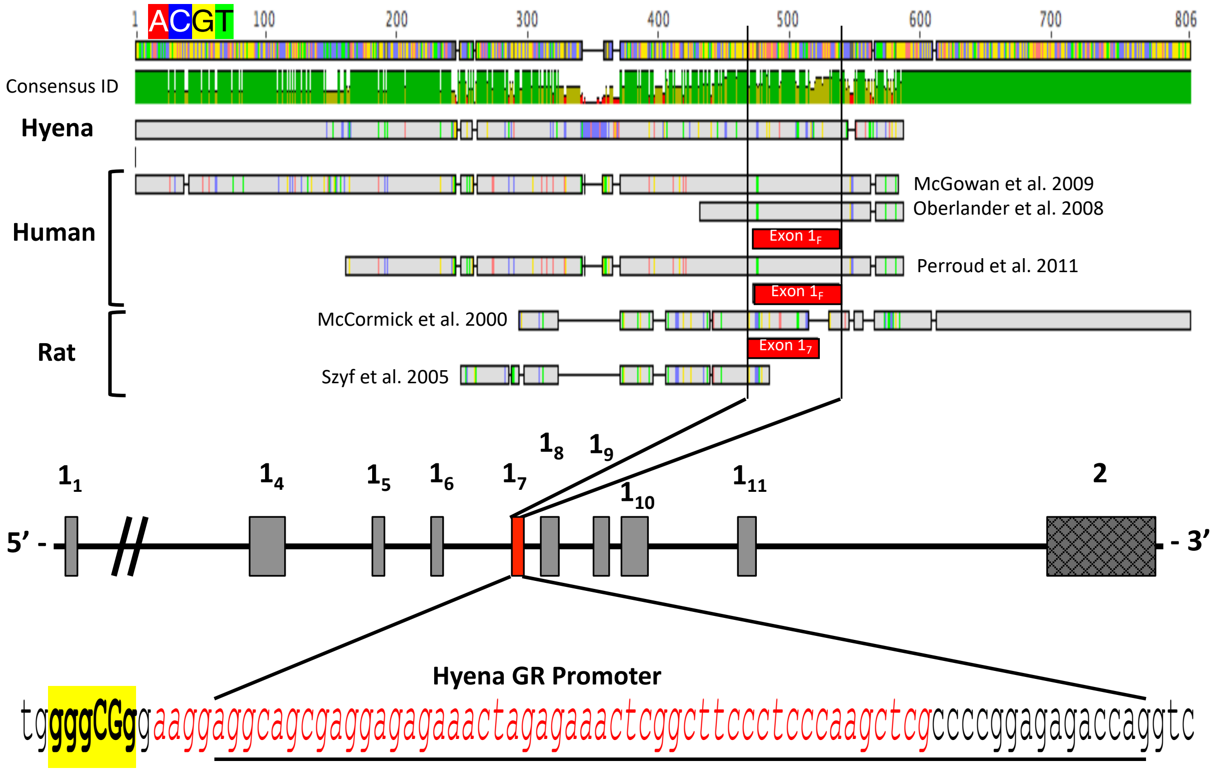


**Supporting Figure 6**. Cross species comparison (spotted hyenas, humans and rats) of the GR promoter region that is the target of gene-specific DNA methylation studies. Base pair consensus from amplicons reported in multiple published studies as well as our spotted hyena amplicon are shown. Here we show hyena DNA sequence that corresponds with human exon $1_F$, underlined and with rat exon $1_7$, red and italicized. Also identified in yellow highlighted and bold text is the putative transcription factor binding site for NGFI-A.

*Demographic, social experience, and ecological covariates*

We estimated each hyena's age with an accuracy ± 7 days based on the behaviors and morphology of each cub when they were first observed outside of their natal den [25]. Once a hyena was three months old, we determined their sex based on the glans morphology of their erect phallus [26]. Maternal rank the year each offspring was born was calculated based on each adult female's wins and losses during agonistic interactions for a given year [27–29]. We normalized adult female ranks each year on a scale of -1 (lowest rank) to 1 (highest rank) to account for changes in group size. We identified if young hyenas had a sibling or not and grouped them as twins or singletons, respectively. Using our detailed demographic data, we recorded each mom's parity for a given offspring and categorized this variable as either primiparous or multiparous. We also recorded clan size over the duration of our study period.

Next, we categorized offspring hyenas into groups according to whether they were born in low, medium, or high human disturbance based on illegal livestock grazing in the park [30]. We also grouped them into two categories of food availability based on the time of year that they were born. Birth dates roughly corresponding to the annual wildebeest and zebra migration from June – November were categorized as migration present and hyenas born December- May were categorized as migration absent.

*Statistical Analyses*

Prior to formal analysis, we assessed the distribution of continuous variables as well as frequency tabulations of categorical variables to check for deviations from normality, errors in the data, missing values, and sample sizes within strata. We constructed boxplots to identify outliers and viewed scatterplots to check linearity of associations between covariates and our outcome variables. We also examined bivariate associations between dependent and independent variables to help identify potential confounding variables that could influence the relationships of interest in our final models.

We organized our analyses into four parts and included two primary data sets for both global and genome-wide DNA methylation. The global DNA methylation data set used in our analyses comprises four overlapping data subsets with information on: (1) DNA methylation measures from 186 cub and subadult hyenas (age ≤24 months), (2) maternal care behaviors from focal animal survey (FAS) data on 258 unique mother-cub pairs, (3) social network data from 115 hyenas during two early phases of development: the communal den (CD) period when a young hyena resided exclusively at the communal den, and the den independent (DI) period, which began when cubs were found away from the communal den on at least 4 consecutive occasions, and (4) fecal Glucocorticoid Metabolites (fGCMs) from 268 adult (≥24 months old) hyenas. The overlap among data sets, and the final analytical samples sizes for analysis parts one through three, are shown in **Supporting Figure 7**. It is also worth noting that the earliest date of the maternal care FAS session and the start date of the social network period preceded the immobilization date when blood was drawn, and DNA methylation was measured. We also included only fGMCs that were obtained after assessments of maternal care behaviors, social network metrics, and %CCGG DNA methylation in order to preserve temporal relationships between our explanatory variables and our outcome variables to improve causal inference.

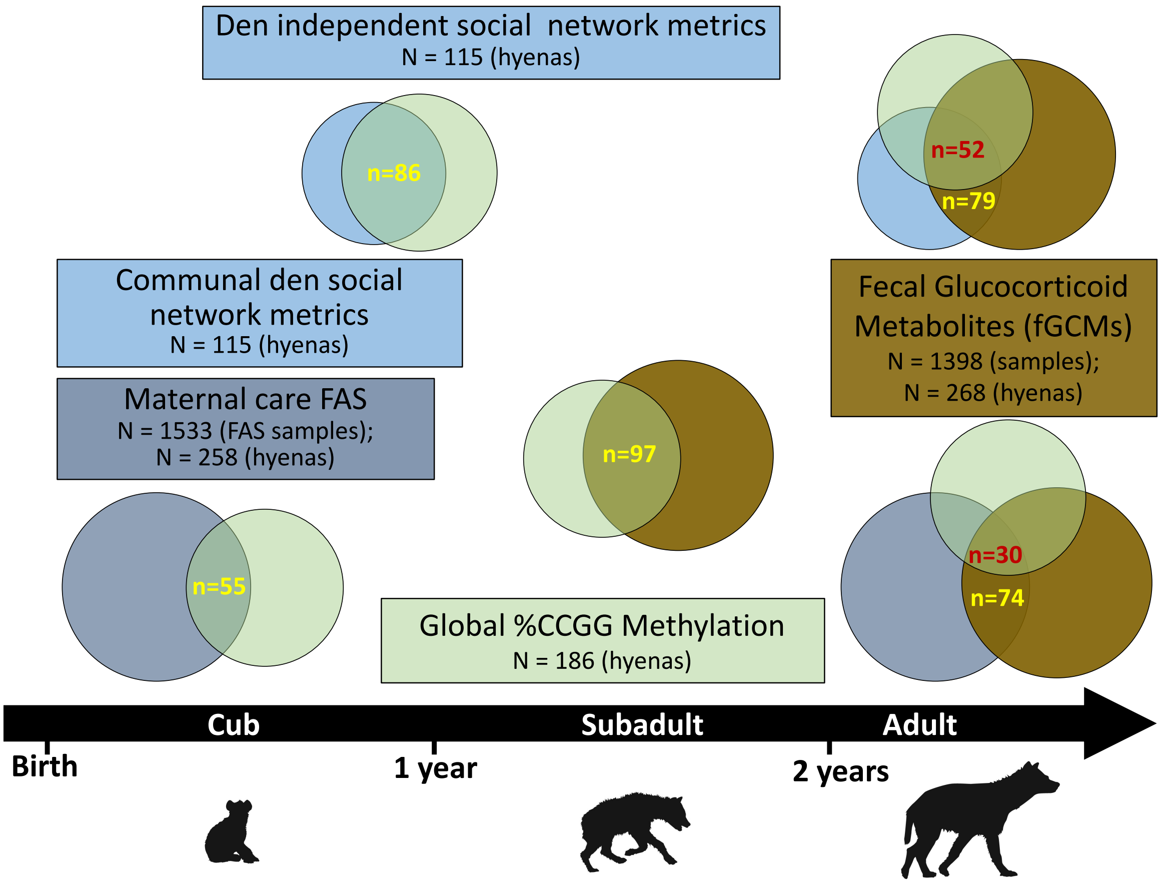


1.

**Supporting Figure 7**. Overview of the sampling design and sample sizes from the four data sets used in the analyses (parts 1-3). The timeline on the bottom shows the different life stages of hyenas. Each color of box represents a distinct type of data collected from our study population, and the edges of boxes roughly correspond to the hyenas' ages when samples were collected. Overall sample sizes and numbers of individual hyenas after data cleaning are indicated inside each box. Venn diagrams show the overlap in sample size among the different data sets and that correspond with the statistical models (yellow = overlap of two data sets, and red = overlap of three data sets for mediation analyses).

The fourth part of our analysis focused on our genome-wide DNA methylation data. Using mERRBS, we assayed 29 hyenas, 25 of which also had fGCMs, and 12 of which additionally had information on maternal rank and maternal care metrics from FAS. Similar to the maternal care metrics, we summarized repeated measures of fGCMs via a mixed-effects linear regression model in which the natural log of fGCMs was the dependent variable. We controlled for each hyena's age in months, reproductive state, and time of day (AM vs PM) when the fecal sample was collected, a random intercept for offspring ID to account for correlations between samples collected from the same individual, and an unstructured covariance matrix. Repeated fGCMs were limited to when hyenas were 1 yr or older so that our assessment of the stress phenotype roughly overlapped or proceeded the assessment of genome-wide DNA methylation. We then calculated BLUPS from this mixed-effects model. Here again, the BLUPs effectively represent each hyena's deviation in fGCMs relative to the population average after accounting for key demographic covariates. This method has been previously used to consolidate repeated measurements of a given variable into a single value per individual without making assumptions about the underlying distribution of the data [31,32]. Linear mixed-models were run using the R package lme4 [33].

### Results

*Analysis parts 1-3 background characteristics and descriptive statistics*

Mother and offspring hyena pairs spent a mean ± SD of the proportion of time they were observed together during FAS sessions in close proximity (0.826 ± 0.099), nursing (0.458 ± 0.064), and grooming (0.074 ± 0.053). During the CD period of development, the mean ± SD of association index social network metrics were 57.45 ± 13.29 for degree centrality, 8.85 ± 2.11 for strength, and 6.50 ± 11.81 for betweenness centrality. During the DI stage, the mean ± SD for degree, strength, and betweenness were 61.93 ± 15.94, 4.18 ± 1.54, and 6.79 ± 6.11, respectively. We estimated a mean ± SD of fGCMs of 104.0 ± 95.9 ng/g for 268 hyenas with least one adult fecal sample (we averaged repeated measure for an individual). Finally, we calculated a mean ± SD of 75.53 ± 3.03 %CCGG methylation for the 186 hyena cubs and subadults in our study population.

Bivariate analysis: Global (%CCGG) DNA methylation data set

In bivariate analyses, we assessed crude associations between potential confounding covariates with each variable of interest, namely maternal care FAS behaviors, association social network metrics and fGCMs. Many of these associations have been published previously [5,34], so we do not interpret or discuss bivariate results, but rather report the model estimates in **Supporting Tables 2-5**. Variables associated with %CCGG methylation have recently been reported [11], so we do not show those results here.

**Supporting Table 2.** Bivariate associations of demographic, early social experience, and ecological covariates with maternal care behavior BLUPs from 258 hyena cubs.

| | β (± SE)[†] | | | | | |
|---|---|---|---|---|---|---|
| | **Close proximity** | ***P*** | **Nursing** | ***P*** | **Grooming** | ***P*** |
| **Biological confounding variables** | | | | | | |
| **Sex** | | | | | | |
| Female | 0.00 (Reference) | | 0.00 (Reference) | | 0.00 (Reference) | |
| Male | -0.01 (± 0.01) | 0.497 | 0.00 (± 0.01) | 0.603 | 0.01 (± 0.01) | 0.070 |
| **Social experience confounding variables** | | | | | | |
| **Number of litter mates** | | | | | | |
| Singleton | 0.00 (Reference) | | 0.00 (Reference) | | 0.00 (Reference) | |
| Twins | **0.04 (± 0.01)** | **0.014** | **0.02 (± 0.01)** | **0.010** | **-0.02 (± 0.01)** | **0.002** |
| **Mother's parity** | | | | | | |
| Primiparous | 0.00 (Reference) | | 0.00 (Reference) | | 0.00 (Reference) | |
| Multiparous | -0.02 (± 0.01) | 0.106 | 0.01 (± 0.01) | 0.141 | 0.00 (± 0.01) | 0.789 |
| **Ecological confounding variables** | | | | | | |
| **Human disturbance (year offspring born)** | | | | | | |
| Low | 0.00 (Reference) | | 0.00 (Reference) | | 0.00 (Reference) | |
| Medium | -0.02 (± 0.02) | 0.125 | -0.01 (± 0.01) | 0.141 | -0.01 (± 0.01) | 0.104 |
| High | **0.05 (± 0.02)** | **0.002** | **0.05 (± 0.01)** | **<0.001** | 0.01 (± 0.01) | 0.408 |
| **Migration status on date of birth** | | | | | | |
| Migration absent | 0.00 (Reference) | | 0.00 (Reference) | | 0.00 (Reference) | |
| Migration present | -0.02 (± 0.01) | 0.220 | **-0.02 (± 0.01)** | **0.003** | -0.01 (± 0.01) | 0.449 |

[†]Estimates represent differences in incident rates or proportions of time spent engaged in a particular behavior in relation to the total time the mother-offspring pair were observed together (n = 55

**Supporting Table 3**. Bivariate associations of demographic, early social experience, and ecological covariates with communal den (CD) social network metrics.

| | β (± SE)† | | | | | |
|---|---|---|---|---|---|---|
| | **Degree†** | ***P*** | **Strength** | ***P*** | **Betweeness** | ***P*** |
| **Biological confounding variables** | | | | | | |
| **Sex** | | | | | | |
| Female | 0.00 (Reference) | | 0.00 (Reference) | | 0.00 (Reference) | |
| Male | 3.98 (± 2.49) | 0.113 | 0.60 (± 0.39) | 0.132 | 3.84 (± 2.21) | 0.085 |
| **Social experience confounding variables** | | | | | | |
| **Number of litter mates** | | | | | | |
| Singleton | 0.00 (Reference) | | 0.00 (Reference) | | 0.00 (Reference) | |
| Twins | -1.39 (± 4.03) | 0.744 | 0.47 (± 0.63) | 0.460 | -3.26 (± 1.89) | 0.087 |
| **Clan size** | | | | | | |
| Number of hyenas in clan | **0.54 (± 0.10)** | **<0.001** | 0.03 (± 0.02) | 0.059 | **0.23 (± 0.10)** | **0.020** |
| **Ecological confounding variables** | | | | | | |
| **Human disturbance (year offspring born)** | | | | | | |
| Low | 0.00 (Reference) | | 0.00 (Reference) | | 0.00 (Reference) | |
| Medium | **-8.57 (± 2.66)** | **<0.001** | -0.37 (± 0.41) | 0.369 | 3.75 (± 2.46) | 0.130 |
| High | **12.41 (± 2.79)** | **<0.001** | **1.82 (± 0.50)** | **<0.001** | -0.07 (± 3.02) | 0.981 |
| **Migration status on date of birth** | | | | | | |
| Migration absent | 0.00 (Reference) | | 0.00 (Reference) | | 0.00 (Reference) | |
| Migration present | **-5.96 (± 2.57)** | **0.022** | **-1.64 (± 0.39)** | **<0.001** | -1.36 (± 2.33) | 0.560 |

†Estimates are based on association index networks from 115 communal den dependent hyeans.

**Supporting Table 4**. Bivariate associations of demographic, early social experience, and ecological covariates with DI social network metrics.

| | β (± SE)† | | | | | |
|---|---|---|---|---|---|---|
| | **Degree** | ***P*** | **Strength** | ***P*** | **Betweeness** | ***P*** |
| **Biological confounding variables** | | | | | | |
| **Sex** | | | | | | |
| Female | 0.00 (Reference) | | 0.00 (Reference) | | 0.00 (Reference) | |
| Male | 2.70 (± 3.01) | 0.371 | 0.28(± 0.29) | 0.332 | 0.17 (± 1.16) | 0.884 |
| **Social experience confounding variables** | | | | | | |
| **Number of litter mates** | | | | | | |
| Singleton | 0.00 (Reference) | | 0.00 (Reference) | | 0.00 (Reference) | |
| Twins | 5.15 (± 4.86) | 0.292 | 0.80 (± 0.46) | 0.086 | -1.30 (± 1.66) | 0.435 |
| **Clan size** | | | | | | |
| Number of hyenas in clan | **0.77 (± 0.08)** | **<0.001** | **0.03 (± 0.01)** | **0.014** | **0.13 (± 0.04)** | **0.002** |
| **Ecological confounding variables** | | | | | | |
| **Human disturbance (year offspring born)** | | | | | | |
| Low | 0.00 (Reference) | | 0.00 (Reference) | | 0.00 (Reference) | |
| Medium | **-12.29 (± 2.59)** | **<0.001** | **-0.93 (± 0.30)** | **0.002** | -1.22 (± 1.28) | 0.045 |
| High | **14.75 (± 3.20)** | **<0.001** | 0.69 (± 0.36) | 0.058 | -0.66 (± 1.58) | 0.676 |
| **Migration status on date of birth** | | | | | | |
| Migration absent | 0.00 (Reference) | | 0.00 (Reference) | | 0.00 (Reference) | |
| Migration present | -0.82 (± 3.15) | 0.796 | 0.33 (± 0.30) | 0.282 | -1.83 (± 1.20) | 0.130 |

†Estimates are based on association index networks from 115 communal den independent hyeans.

**Supporting Table 5.** Bivariate associations of sample collection variables and adult fecal Glucocorticoid Metabolites (fGCMs).

| | 0 | β (± SE)† | |
|---|---|---|---|
| **Potential confounding variables** | | **fGCMs** | ***P*** |
| **Sex** | | | |
| Female | | 0.00 (Reference) | |
| Male | | **-0.72 (± 0.10)** | **<0.001** |
| **Standardized Maternal Rank (yr offspring born)** | | -0.13 (± 0.09) | 0.150 |
| **Human disturbance (yr offspring born)** | | | |
| Low | | 0.00 (Reference) | |
| Medium | | **0.24 (± 0.08)** | **0.004** |
| High | | -0.08 (± 0.09) | 0.366 |
| **Precision covariates (assessed on fecal collection date)** | | | |
| **Age (months)** | | **0.005 (± 0.001)** | **<0.001** |
| **Reproductive state** | | | |
| Nulliparous | | 0.00 (Reference) | |
| Pregnant | | **0.98(± 0.11)** | **<0.001** |
| Lactating | | **0.59 (± 0.09)** | **<0.001** |
| Other | | **0.64 (± 0.11)** | **<0.001** |
| Male | | -0.18 (± 0.12) | 0.130 |
| **Time of day** | | | |
| AM | | 0.00 (Reference) | |
| PM | | **-0.50 (± 0.05)** | **<0.001** |
| **Migration status** | | | |
| Migration absent | | 0.00 (Reference) | |
| Migration present | | 0.09 (± 0.06) | 0.091 |

†Beta estimate are differences in each categorical variable from the reference group in fGCMs (ng/g) on the natural log scale from mixed models in which hyena ID was included as a random intercept. Models include 1398 fGCMs samples collected from 268 adult (≥24 months old) hyenas.

*Mediation analyses*

**Supporting Table 6**. Association of maternal care and early life social network metrics with %CCGG methylation and adult fecal Glucocorticoid Metabolites (fGCMs). Summary of results from steps 1-2 of a mediation analysis. X represents the explanatory variable (early life social experience), M represents the potential mediator (%CCGG methylation), and Y represents the outcome (fGCMs).

| | | β (95% CI) | | | |
|---|---|---|---|---|---|
| | ***N*** | **Step 1: X → Y †** | **Step 2a: X → M ‡** | | **Step 2b: M → Y †** |
| **Maternal Care FAS (per 1-SD)** | | | | **DNA Methylation (per 1-SD) §** | |
| Close proximity | 30 | -0.08 (-0.38, 0.21) | **1.45 (0.63, 2.27)** | %CCGG | -0.02 (-0.36, 0.33) |
| Nursing | 30 | -0.07 (-0.43, 0.31) | 0.53 (-0.73, 1.83) | | |
| Grooming | 30 | -0.16 (-0.43, 0.11) | 0.00 (-0.46, 2.07) | | |
| **CD period (per 1-SD)** | | | | | |
| Degree | 52 | -0.08 (-0.24, 0.09) | 0.17 (-0.87, 1.20) | | |
| Strength | 52 | 0.01 (-0.15, 0.18) | 0.31 (-0.61, 1.23) | | |
| Betweenness | 52 | -0.10 (-0.33, 0.13) | -0.13 (-0.91, 0.66) | **DNA Methylation (per 1-SD) Φ** | |
| **DI period (per 1-SD)** | | | | %CCGG | 0.06 (-0.13, 0.24) |
| Degree | 52 | -0.11 (-0.27, 0.06) | 0.18 (-1.29, 1.64) | | |
| Strength | 52 | -0.08 (-0.24, 0.08) | 0.61 (-0.33, 1.54) | | |
| Betweenness | 52 | -0.06 (-0.22, 0.10) | -0.16 (-1.10, 0.68) | | |

†Beta estimate are fGCMs (ng/g) on the natural log scale from mixed models in which hyena ID was included as a random intercept. Models are adjusted for hyena's age (months), sex, reproductive state among females, and the time of day when the fecal sample was collected.

‡ Models are adjusted for hyena's age when blood was drawn (months), sex, number of littermates, maternal rank, mom's parity / clan size, human disturbance in the birth yr, and the migration status on the birth date.

§ Assocation is based on samples for which there are measures of maternal care, DNA methylation, and fGCMs.

Φ Assocation is based on samples for which there are measures of social connectedness during the CD and DI periods, DNA methylation, and fGCMs.

*Genome-wide (mERRBS) DNA methylation analyses*

We analyzed data from 29 female hyenas which had an average ± SD age of 18.8 ± 3.9 months on the date their DNA methylation was assessed. After processing the data through the bioinformatic pipeline, we quantified DNA methylation at 2,246,867 unique CpG sites that had a minimum of 10x coverage in all 29 hyenas. The average bisulfite conversion rate was 99.4% (SD ± 0.1%) and the average alignment efficiency of reads to the draft spotted hyena genome was 75.5% (SD ± 2.6) for the 29 hyena samples. Of the 29 hyenas for which we had mERRBS data, we also had fecal samples from 25 of them (median of 3 samples per hyena). Mean ± SD later life corticosterone concentration in our study population was 72.9 ± 62.8 ng/g. During the FAS data collection, we observed mother-offspring pairs together on average for 26 minutes per FAS session, with a median of 2.5 sessions recorded per pair. In this subsample, mothers spent an average of 79.8% (SD ± 30.0%) of their time in close proximity to offspring, 66.4% (SD ± 31.2%) of time nursing, and 4.9% (± 7.3%) of time grooming.

The uncorrected EWAS revealed 97,745 stress related differentially methylated CpG sites. The distribution of the observed p-values generated from the binomial regression based EWAS did not indicate any substantial outliers and generally conformed with the expected p-value distribution (**Supporting Figure 8, a**). Additionally, the p-values from the EWAS generally followed a uniform distribution (**Supporting Figure 8, b**).

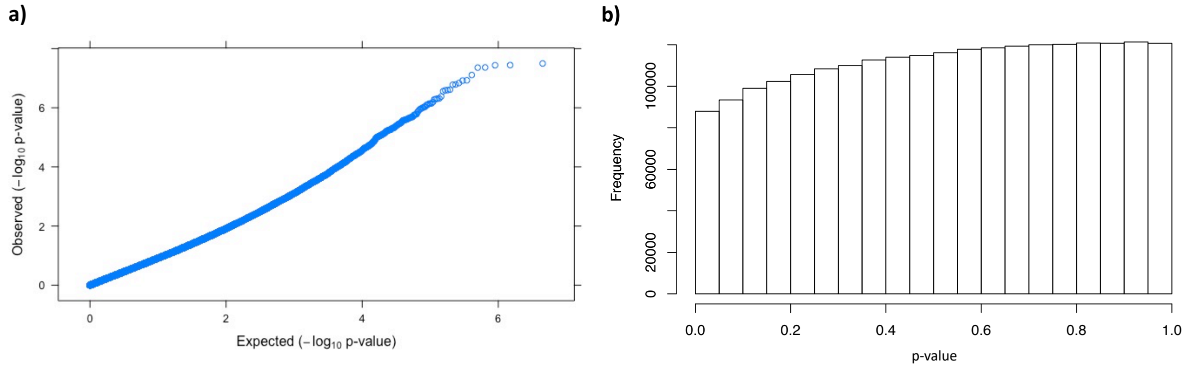


**Supporting Figure 8**. Diagnostic plots for stress related EWAS. **a)** A QQ-plot showing the observed $-\log_{10}$ p-values versus the expected $-\log_{10}$ p-values. **b)** A frequency histogram showing a relatively uniform distribution of p-values generated from the EWAS.

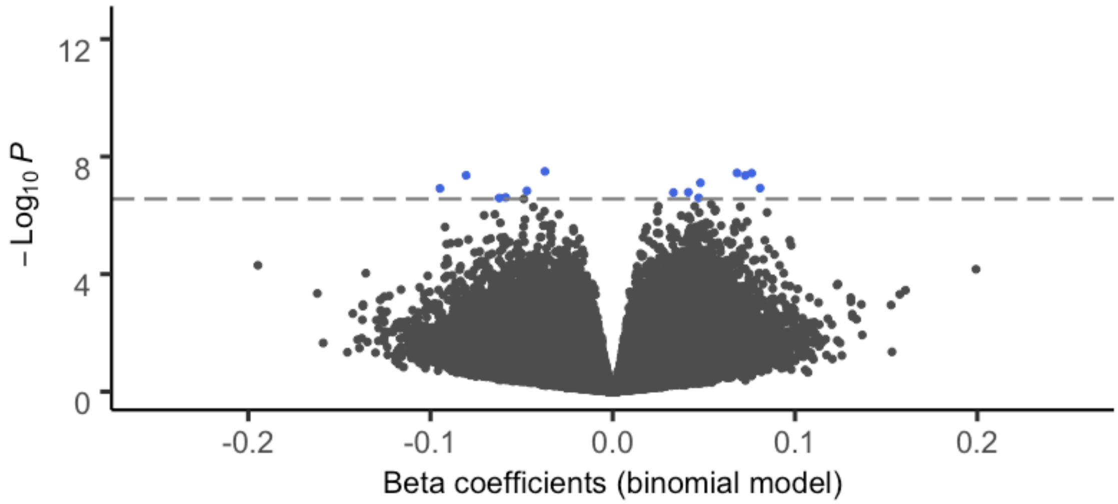


**Supporting Figure 9**. Volcano plot of the $-\log_{10}$ p-values by the beta coefficients from a binomial regression epigenome wide association study (EWAS). Counts of methylated over total cytosines from enhanced reduced representation bisulfite sequencing data were modeled as a function of variation in subadult and adult spotted hyena fecal Glucocorticoid Metabolites (fGCMs). The horizontal line represents the Benjamini-Hochberg false discovery rate (FDR) cutoff of 0.05%, above which the blue points are significant.

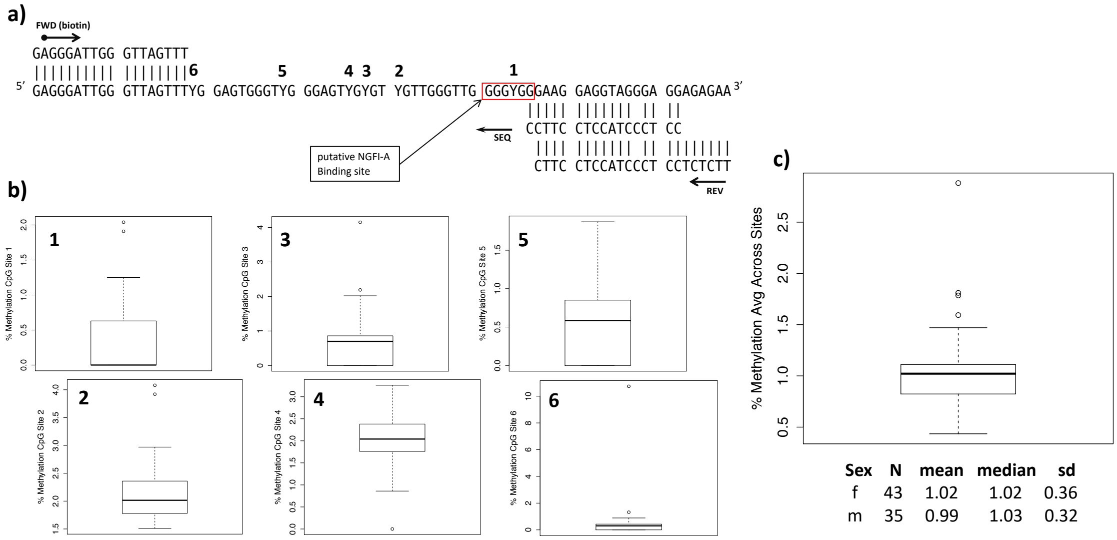


| Sex | N | mean | median | sd |
|---|---|---|---|---|
| f | 43 | 1.02 | 1.02 | 0.36 |
| m | 35 | 0.99 | 1.03 | 0.32 |

**Supporting Figure 10**. Quantitative measurement of DNA methylation at 6 CpG sites in the hyena putative GR promoter in a DNA sequence that was targeted in previous rat and human studies. **a)** Target DNA sequence with potentially methylated cytosine bases identified with a 'Y' and numbered according to pyrosequencing order. Putative transcription factor binding site is identified in the red box and forward, reverse and sequencing primers are shown on the DNA strand. **b)** Boxplots showing site-specific CpG percent methylation. **c)** Boxplot and descriptive statistics showing percent methylation values averaged across all 6 CpG sites from 78 hyenas (note some samples did not pass the pyrosequencing quality standards, so the total samples size dropped from 96 hyenas).
